# Unsupervised 3D classification and characterisation of osteocyte and chondrocyte lacunae in the murine osteochondral unit

**DOI:** 10.64898/2026.10.07.757361

**Authors:** Anik Banerjee, Pholpat Durongbhan, Piyush Uniyal, Valentine Gaillard, Jemima E Schadow, Emma C Boersma, Kathryn S. Stok, Davide Ruffoni, G Harry van Lenthe

## Abstract

In the murine osteochondral unit, morphology of chondrocyte lacunae (CCL) within calcified articular cartilage and osteocyte lacunae (OCL) within the underlying subchondral bone reflects local biomechanical conditions that are altered in disease, making reliable discrimination of both lacunar populations important for studying osteochondral physiology and pathology. Histology is generally considered the gold standard for identifying CCL and OCL. However, it is inherently limited to two-dimensional assessment and a relatively small tissue section, restricting comprehensive characterisation of three-dimensional lacunar structure. Synchrotron-radiation micro-computed tomography (SR-microCT) enables volumetric quantification of lacunae; where classification is typically based on the position of each lacuna relative to the cement line separating calcified cartilage from bone, requiring manual delineation of this convoluted interface. Such approaches are labour-intensive, operator-dependent, and rely on access to specialised synchrotron infrastructure that is not widely available. To address these limitations, we present a fully automated, annotation-free pipeline for classifying OCL and CCL in the mouse osteochondral unit based on desktop micro-computed tomography (microCT), independent of cement line identification. The proposed unsupervised workflow requires neither tissue separation nor labelled training data, enabling objective and scalable discrimination of lacunar populations directly from volumetric image data. Seven morphometric and positional features per lacuna were extracted from SR-microCT and desktop microCT datasets and analysed using Uniform Manifold Approximation and Projection (UMAP). Validation of the pipeline outcome on SR-microCT against manual classification across specimens yielded very-high agreement, with Gwet’s AC1, the Matthews correlation coefficient, and balanced accuracy each approaching 0.99, in both healthy and post-traumatic osteoarthritis specimens. Such a workflow can be readily adapted for the investigation of osteochondral interface in various other species.

## INTRODUCTION

The morphology of osteocyte lacunae (OCL) in bone tissue and chondrocyte lacunae (CCL) in calcified articular cartilage tissue is influenced by the local biomechanical environment. Cell morphology is also believed to have a role in the sensing and transduction of mechanical signals [1], [2]. Strong modifications in OCL and CCL morphologies have been linked to decreased bone mechanoresponsiveness and to the onset and progression of osteoarthritis [3], [4], [5]. These morphological changes are particularly pronounced in the mineralised tissue of the osteochondral unit of the proximal tibia, which comprises calcified articular cartilage and subchondral bone. These two tissues, separated by the osteochondral cement line, differ in mineral content, mechanical and biological [6], [7].

Lacunar size, shape, and spatial distribution are quantified from 2D sections [8] or 3D imaging to compare populations across age, disease grade, and treatment conditions [2], [9], and interpreting these measurements requires distinguishing OCL from CCL. Therefore, a common approach to the study of lacunar morphology starts with the separation of calcified articular cartilage and subchondral bone by identifying the osteochondral cement line delineating the interface between the two mineralised tissues. Histological techniques [10] and quantitative backscattered electron imaging [11] can be used to separate calcified articular cartilage from subchondral bone by exploiting differences in biochemical composition and mineral content. However, the planar nature of these techniques limits morphometric extraction to 2D. High-resolution volumetric imaging techniques, such as conventional absorption-based desktop micro-computed tomography (microCT) and synchrotron radiation phase-contrast microCT (SR-microCT), provide full 3D lacunar morphometry across the imaged tissue volume.

Desktop microCT is more widely accessible and cost-effective than SR-microCT yet cannot clearly resolve the osteochondral cement line [12]. SR-microCT produces higher signal-to-noise ratios sufficient to delineate the cement line directly [13], although existing SR-microCT workflows still depend on manual tracing to reconstruct it. Finnilä et al., manually segmented the subchondral bone and articular cartilage in human sample using both histology sections and microCT slices [14]. A commercial GUI-based software was used to generate binary masks and delineating the calcified cartilage and subchondral bone compartments in murine tibia, requiring human supervision[15]. Comparable challenges have been reported in other species: deep-learning approaches have demonstrated the feasibility of automated tissue segmentation in rabbit knee microCT [16] and in human, lacunar quantification at the osteochondral interface with SR-microCT also required labelled training data [13]. In another study, the interface was identified based on the morphological features and contrast differences and delineated manually[17]. No existing workflow therefore classifies osteocyte and chondrocyte lacunae in the murine osteochondral unit without manual tissue separation, cement line tracing, or labelled training data. At the population level OCL and CCL are morphologically distinct. OCL are smaller and more elongated [9], [18], whereas CCL are larger, more spherical, and may be organised into chondrons or exhibit hypertrophy [19], [20]. However, at the level of individual lacunae, the morphological parameter ranges of OCL and CCL overlap substantially. Therefore, no single criterion, whether volume, shape, or orientation, can reliably differentiate them [13], [15].

Against this background, the present study pursued two primary aims. First, to develop a fully automated, annotation-free 3D pipeline for OCL and CCL classification in SR-microCT data and to validate it against manual classification. Second, to demonstrate cross-modality equivalence between SR-microCT and microCT, establishing indirect validation of the developed methodology when using microCT data. Transferability to 2D planar dataset was explored as a supplementary analysis.

## METHODS

### A. Validating the pipeline on SR-microCT

#### 1. Samples and image acquisition

All animal protocols were approved by the University of Melbourne animal ethics committee (#24339) and conducted in compliance with the Australian code for the care and use of animals for scientific purposes. Samples obtained from the University of Liège were collected in the framework of an organ donation programme(ethical approval: ULg-IACUC-23–2580).

Six male C57Bl/6 mice were used for SR-microCT imaging. Three baseline control animals were sacrificed at 10 weeks of age without surgical intervention: two obtained from the University of Liège and one purchased from Ozgene ARC (Bentley, Australia) and housed at the University of Melbourne under climate-controlled conditions. The remaining three animals, also purchased from Ozgene ARC and housed at the University of Melbourne, underwent destabilisation of the medial meniscus (DMM) on the right knee at 10 weeks of age and were sacrificed 56 days post-surgery (18 weeks of age). Following sacrifice, the right hind limb was harvested, wrapped in PBS-soaked gauze, and stored at −20°C until processing. Limbs were then cleaned of soft tissue, fixed in 70% ethanol for 24 hours, and dehydrated through graded ethanol (80%, 90%, 100%; 24 hours per step), followed by acetone immersion to enhance radiographic contrast between mineralised and non-mineralised tissue [21].

SR-microCT imaging of the medial plateau was performed at the ANATOMIX beamline at Synchrotron SOLEIL, France, in propagation-based phase-contrast mode. A parallel beam with an estimated peak energy of approximately 31.5 keV was used in combination with a CMOS ORCA-Flash4 detector. Depending on the sample, 4700–5900 projections were acquired over a 360° rotation, with an exposure of 40 ms per projection, giving a total duration of about 7 min per sample. SR propagation-based phase-contrast micro-CT images were reconstructed with Paganin phase retrieval and filtered back projection using the PyHST2 software (version 2021c) developed at ESRF [22]. The 3D reconstructed volumes consisted of voxels with an isotropic voxel size of 0.65 µm., Samples were positioned vertically and the entire osteochondral region, including calcified cartilage and subchondral bone, was scanned. For the analysis, a volume of interest, positioned midway through the medial plateau, and having dimensions of 800 × 400 × 420 μm³ was considered (*Fig.1*).

**Fig. 1.**
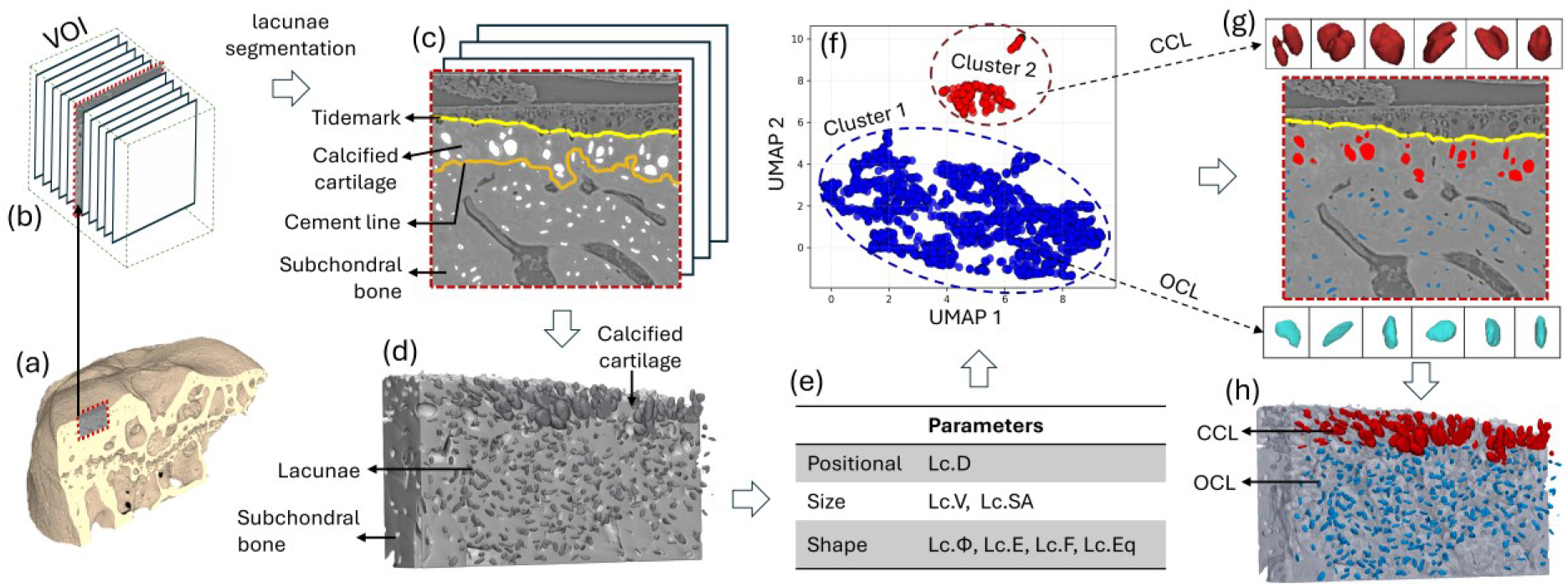
Overview of the unsupervised lacunae classification pipeline. (a) Illustration of the medial tibial plateau of a murine proximal tibia imaged by synchrotron-radiation micro-computed tomography (SR-microCT), (b) A single slice extracted from within the VOI, showing subchondral bone (c) segmented lacunae from individual slices (d) 3D rendering of segmented lacunae, shown against the subchondral bone in the background(e) seven descriptors were extracted: a positional parameter (Lc.D, distance from the outer mineralised surface), two size parameters (Lc.V, volume; Lc.SA, surface area), and four shape parameters (Lc.φ, sphericity; Lc.E, elongation; Lc.F, flatness; Lc.Eq, equancy). (f) Uniform manifold approximation and projection (UMAP) with hierarchical density-based clustering (HDBSCAN) separated the lacunae into two populations, identified as calcified chondrocyte lacunae (CCL, red), rounded and located near the joint surface, and osteocyte lacunae (OCL, blue), (g) illustration of segmented chondrocytes and osteocytes after volume rendering and cluster labels were back-projected onto the original image stack, (h) segmented CCL and OCL displayed on the same VOL, with subchondral bone in the background.

#### 2. Image Processing and Feature Extraction

A 3D Gaussian filter (σ = 0.8) was applied to improve the signal-to-noise ratio. Lacunae were then segmented from the surrounding mineralised matrix (comprising both calcified articular cartilage and subchondral bone) using Otsu automatic thresholding, yielding a binary image of pore spaces. Image processing and morphological parameters for each lacuna were calculated using BoneJ (version 1.54p, NIH, USA). Lacunae with volumes smaller than 100 µm³ and larger than 80,000 µm³ were excluded to eliminate noise artefacts and large non-lacunar voids, respectively, consistent with prior literature on OCL and CCL morphometry [2]. Vascular channels, identified as predominantly large, connected pores, were removed using the “Remove Largest Volume” command in the MorphoLibJ plugin [23]. Remaining non-lacunar artefacts (e.g., cracks) were excluded by retaining only porosities with a ratio of largest –to-shortest eigenvalue below 25. All partially rendered lacunae at the edges of the volume of interest (VOI) were excluded.

Six morphological parameters were extracted per lacuna using BoneJ, as summarised in *Table 1*. A seventh positional parameter, lacunar distance (Lc.D), was additionally computed for each lacuna. Lc.D is the distance between the lacuna and the tidemark. the outer boundary of the segmented volume is assumed as the tidemark. Lacunae with Lc.D values between 0 and 10 µm were excluded to eliminate edge artefacts and partially segmented structures near the tidemark. Lc.D is further discussed in *Supplementary Methods A3 (Fig. S1)*.

**Table 1.**
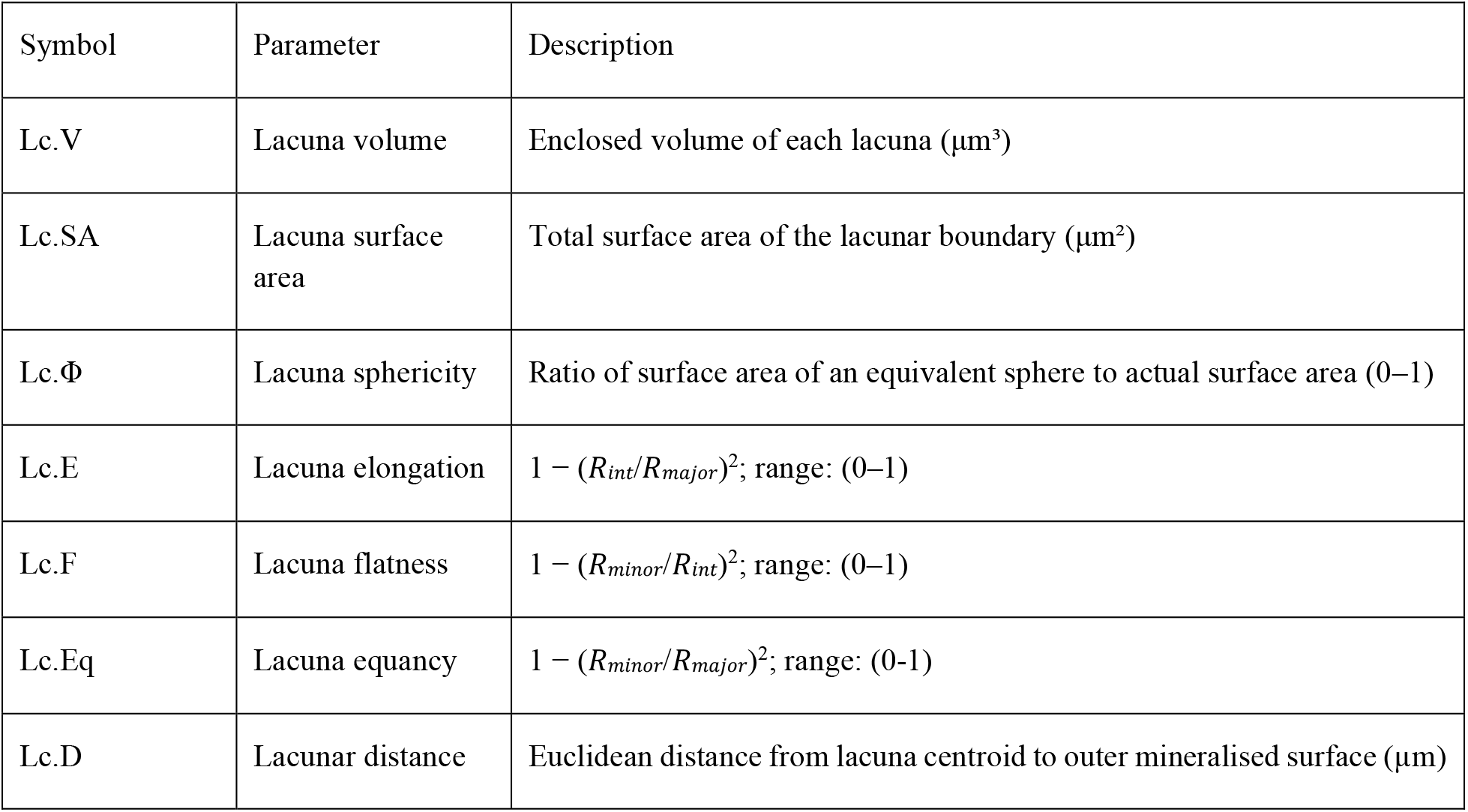
Morphometric and positional parameters extracted per lacuna in the 3D pipeline.

| Symbol | Parameter | Description |
| --- | --- | --- |
| Lc.V | Lacuna volume | Enclosed volume of each lacuna ($\mu\text{m}^3$) |
| Lc.SA | Lacuna surface area | Total surface area of the lacunar boundary ($\mu\text{m}^2$) |
| Lc.$\Phi$ | Lacuna sphericity | Ratio of surface area of an equivalent sphere to actual surface area (0–1) |
| Lc.E | Lacuna elongation | $1 - (R_{\text{int}}/R_{\text{major}})^2$; range: (0–1) |
| Lc.F | Lacuna flatness | $1 - (R_{\text{minor}}/R_{\text{int}})^2$; range: (0–1) |
| Lc.Eq | Lacuna equancy | $1 - (R_{\text{minor}}/R_{\text{major}})^2$; range: (0–1) |
| Lc.D | Lacunar distance | Euclidean distance from lacuna centroid to outer mineralised surface ($\mu\text{m}$) |

The classification framework was adapted for two-dimensional imaging to assess transferability beyond volumetric modalities. Adapted 2D parameters and pipeline details are described in *Supplementary Methods (B.1–B.5* and *Table S1*).

#### 3. UMAP clustering

UMAP was applied across a hyperparameter sweep of *n_neighbors* ∈ {5, 10, 20, 30, 40, 50, 60, 70, 80, 90, 100} and *min_dist* ∈ {0.01, 0.05, 0.10, 0.15, 0.20}, with *n_components* = 3 and *random_state* at scikit-learn default, yielding 55 configurations per dataset. HDBSCAN was subsequently applied to the UMAP embedding to identify clusters without any predefined cluster count. Silhouette score heatmaps were generated across 55 UMAP hyperparameter configurations (*n_neighbors*: 5–100; *min_dist*: 0.01–0.20) to identify the pair of hyperparameters that yielded the best cluster separation for each 3D modality and displayed as a heatmap, where a score of 0 indicates no meaningful cluster separation and indicates perfect separation [24]. The configuration with the highest silhouette score lead to two identifiable clusters for selection (*Fig. 2*).

**Fig. 2.**
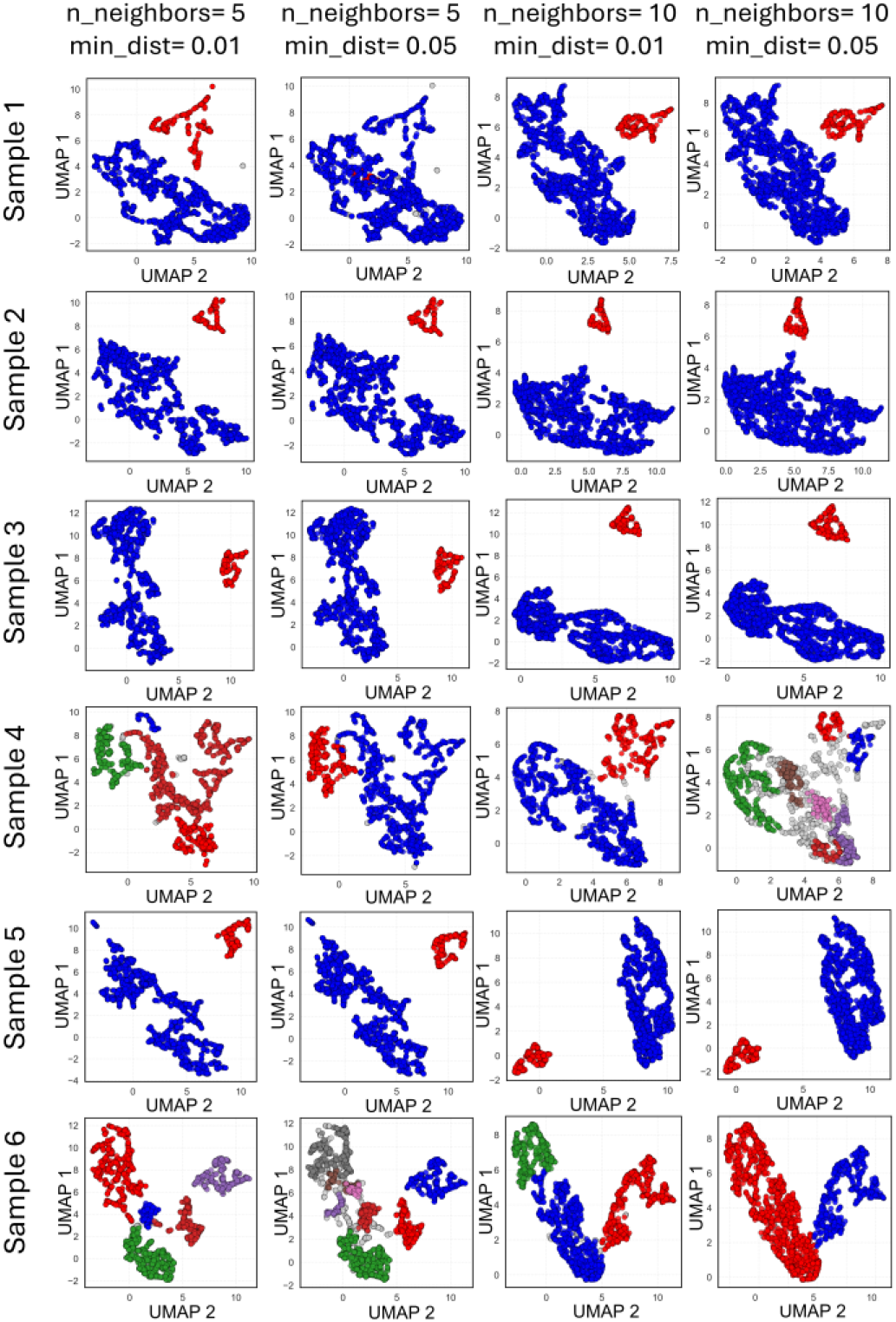
UMAP embeddings under a fixed hyperparameter configuration. Each column corresponds to four fixed hyperparameter combinations (n_neighbors = 5, min_dist = 0.01; n_neighbors = 5, min_dist = 0.05; n_neighbors = 10, min_dist = 0.01; n_neighbors = 10, min_dist = 0.05) applied across all six SR-microCT specimens. Each row corresponds to one specimen (Samples 1–6). Each cluster is represented as a unique colour.

Two other dimensionality reduction techniques, principal Component Analysis (PCA) and t-Distributed Stochastic Neighbour Embedding (t-SNE) were also evaluated to identify the approach best suited to separate OCL and CCL populations. A detailed description can be found in *Supplementary Methods A.4*.

#### 4. Feature-importance analyses

To assess the contribution of each morphological and positional feature to the UMAP embedding and lacuna classification, three analyses were performed: permutation-based feature importance, Spearman rank correlation, and Shapley Additive explanations (SHAP) value analysis. In this study “feature” refers to inputs of the machine-learning stage (permutation importance, SHAP), while “parameter” refers to the underlying morphometric measurements (Lc.V, Lc.SA, etc.).

Permutation-based feature importance was computed for each UMAP component, and the component-level importance scores were averaged to yield a single mean importance score per feature per sample, reported as mean ± standard deviation across samples. A higher score reflects greater contribution of that feature to the classification.[25], [26].

The Spearman rank correlation coefficient was computed between each input parameter and each UMAP component across all lacunae in each sample and visualised as a heatmap. Values range from −1 to +1, where the sign indicates the direction of association and magnitude indicates the degree to which a parameter organises a given UMAP axis [27].

Shapley Additive explanations (SHAP) value analysis was applied at the individual lacuna level [28]. The Spearman rank correlation coefficient (ρ) between each parameter’s raw value and its SHAP contribution was computed per modality and displayed as a horizontal diverging bar chart. Bars extending right (positive ρ) indicate CCL tendency and bars extending left (negative ρ) indicate OCL tendency, with bar length reflecting the magnitude of ρ.

#### 5. Validation against manual ground truth (SR-microCT)

For SR-microCT, the cement line is directly traceable, which allowed manual classification of OCL and CCL. Two image slices per sample were selected from the stacks. A trained observer (AB) manually classified each lacuna as OCL or CCL based on its position relative to the visible cement line. These manual counts served as ground truth against which pipeline-generated labels were compared. A total of 12 SR-microCT slices from six samples were analysed to evaluate pipeline performance against manual classification. Across these 12 slices, 1,294 lacunae were manually classified (mean 107.9 ± 26.4 per slice; OCL 73.7 ± 26.2, CCL 34.2 ± 15.7).

#### 6. Performance metrics

Pipeline performance was evaluated against manual classification using a confusion matrix framework, true positives (TP: OCL correctly classified), true negatives (TN: CCL correctly classified), false positives (FP: CCL incorrectly classified as OCL), and false negatives (FN: OCL incorrectly classified as CCL). Three metrics are reported: Gwet’s AC1 for inter-rater agreement [29], Matthews Correlation Coefficient (MCC) [30], [31], and balanced accuracy. Full equations and interpretation criteria for each of these metrics are provided in *Supplementary Methods A.6*.

### B. Validating the pipeline on microCT

#### 1. Samples and Image Acquisition

After establishing and validating the pipeline on microCT data, we tested the performance of the pipeline with microCT-based data of knees from six C57BL/6 male mice. The dataset included three PTOA samples and three healthy control samples. Only the healthy samples (n=3) were scanned in both modalities.

MicroCT scanning was performed on a TESCAN UniTOM HR device (Tescan, Brno, Czech Republic) at a voxel size of 0.7 µm. Scans were acquired with an exposure time of 650 ms, averaging 5 frames per projection, with 3000 equispaced projections over a 360° rotation. The X-ray source was set to nanofocus mode ((power= 0.8 W; tube voltage= 70kV).. Volumes were reconstructed using filtered back-projection algorithm in Panthera™ (Tescan proprietary software, version 1.3.1)Reconstruction parameters included ring-artefact reduction, beam hardening correction (BHC = 0.1), and a sample-specific centre of rotation (COR) adjustment.

#### 2. Image processing and feature extraction

Image processing and feature extraction followed the exact same procedure as described before for the SR-microCT images (*Methods A.2*).

#### 3. Feature-importance analyses

Feature-importance analyses followed the exact same procedure as described for SR-microCT.

#### 4. Cross-modality validation

For microCT, the cement line is not visible, and therefore direct manual classification is not feasible. Validation was therefore performed by a cross-modality comparison: the full pipeline was independently applied to microCT data and to the corresponding SR-microCT datasets from the same three specimens, covering the same anatomical region selected using identical anatomical landmarks (medial plateau midline in the coronal plane). Pipeline outputs were compared across various metrics: UMAP embedding structure; permutation-based feature importance; Spearman rank correlation; SHAP analysis; mean silhouette score; and cluster count. When the two modalities produce equivalent classification across the selected metrics, the microCT pipeline can be considered to perform equally well as the SR-microCT pipeline, even without a visible cement line. Agreement between modalities constitutes indirect validation of the microCT pipeline.

### C. 2D pipeline

The classification framework was adapted for two-dimensional imaging to assess transferability beyond volumetric CT modalities. It was applied to SR-microCT cross-sections and to histological sections from an independent cohort originally prepared as part of a previous study [32]. Adapted 2D morphometric parameters, sample preparation, and pipeline details are described in *Supplementary Methods B.1–B.5* and *Table S1*.

#### Statistics and software

All statistical analyses were performed in Python 3.10 using scikit-learn (v1.2.0)[33], UMAP-learn[34], and HDBSCAN. Data handling used pandas and NumPy. Visualisations were generated using Matplotlib (v3.6.2), Seaborn (v0.12.2), and Plotly (v5.11.0). Random seeds were fixed for reproducibility: UMAP used the scikit-learn default *random_state*; t-SNE (used only in the Supplementary analyses) used *random_state* = 200. HDBSCAN is deterministic given its inputs. The full pipeline implementation is publicly available at https://gitlab.kuleuven.be/u0161341/uloc.

## RESULTS

### A. Validating the pipeline on SR-microCT

#### 1. UMAP Clustering

UMAP generated well-separated clusters at *n_neighbors* = 5, and as it increased, it failed to form a distinct, compact group with clear boundaries. With decreasing “*min_dist*” the points in each cluster came closer. These results were also consistent with the silhouette scores and number of clusters formed (*Fig. 3*).

**Fig. 3.**
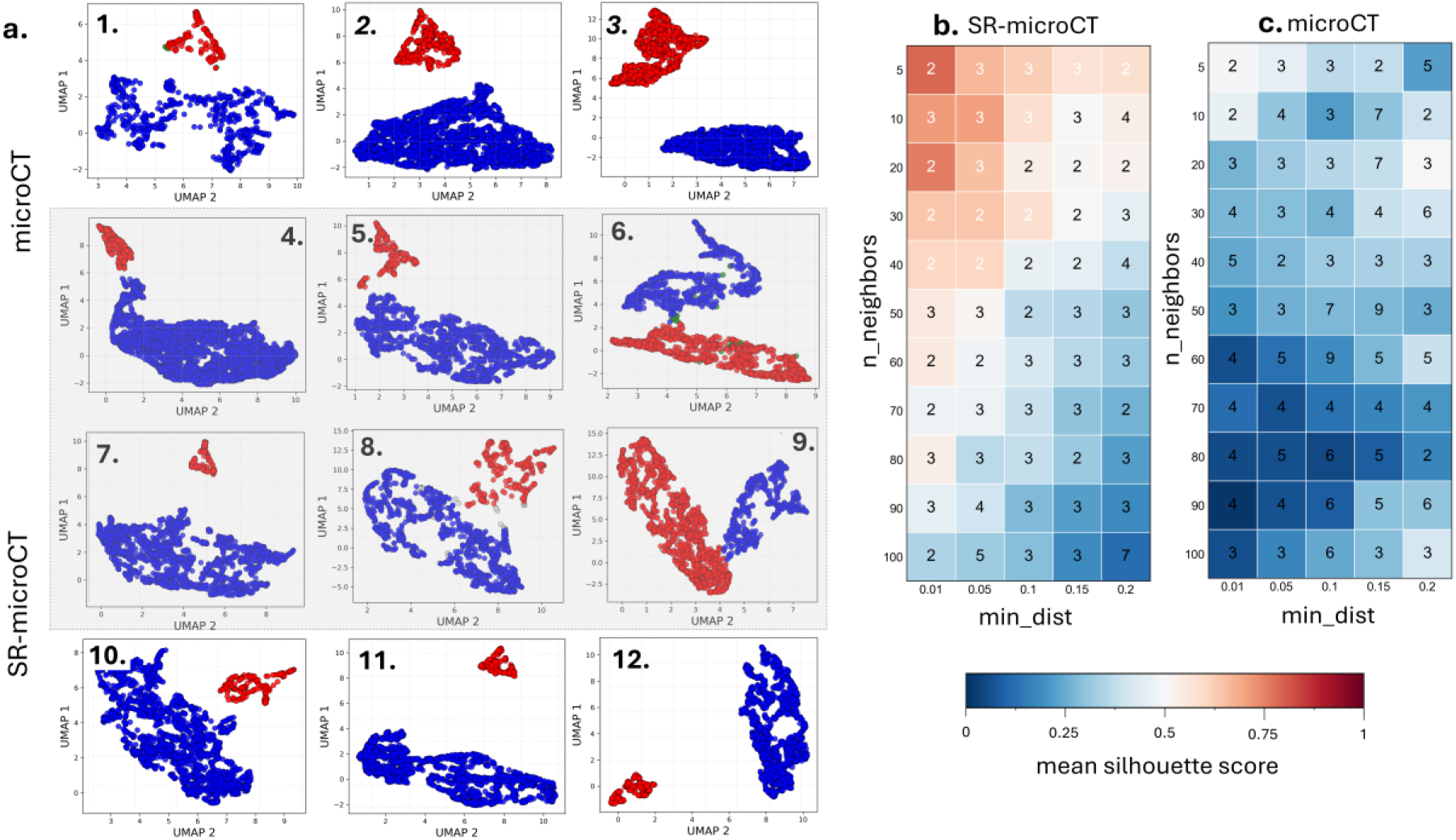
Per-specimen UMAP embeddings and silhouette-score heatmaps for the 3D pipeline. (a) UMAP embeddings (UMAP 1 vs UMAP 2) are shown for microCT specimens (1–6) in the top and SR-microCT specimens (7-12) in the bottom, with cluster 1 in red and cluster 2 in blue. (4 and 7), (5 and 8), and (6 and 9) belong to the same mice (similar region in subchondral bone); (b) Silhouette-score heatmap for SR-microCT and (c) for microCT, spanning n_neighbors (5–100, vertical axis) and min_dist (0.01– 0.20, horizontal axis); each cell is annotated with the number of clusters returned by HDBSCAN and coloured by mean silhouette score from 0 (blue) to 1 (red).

In SR-microCT, the highest mean silhouette scores were recorded as 0.79, with HDBSCAN identifying two clusters in each case. In microCT, silhouette scores were lower across the entire hyperparameter grid; the highest mean score (0.52) was recorded at *n_neighbors* = 5 and *min_dist* = 0.01, again yielding two clusters. The 3D pipeline consistently identified one OCL cluster and one CCL cluster across all specimens and modalities. Back-projection of cluster labels onto the original image stacks confirmed the anatomical identity of each cluster relative to their position on either side of the cement line (*Fig. 1*).

#### 2. Feature importance analysis

##### 2.1. Permutation-Based Feature Importance

Lc.D was the dominant parameter across all 3D datasets, ranking first in the majority of samples. However, large inter-specimen variability was evident: the standard deviation of Lc.D approached or exceeded 50% of its mean in every dataset (SR-microCT: 0.78 ± 0.39; microCT: 0.60 ± 0.38), indicating that feature importance was not consistently stable across individuals. Secondary contributors differed between modalities. In SR-microCT, Lc.V ranked second (0.24 ± 0.16), whereas in microCT, Lc.SA rose to second position (0.47 ± 0.40) with comparable variability. This modality-dependent shift was the most striking difference in parameter hierarchies between the two imaging approaches. Shape parameters (Lc.E, Lc.F, Lc.φ) contributed minimally and consistently across all datasets, scoring below 0.18. Lc.Eq was universally the weakest predictor (mean importance ≤ 0.02) (*Fig. 4f*).

**Fig. 4:**
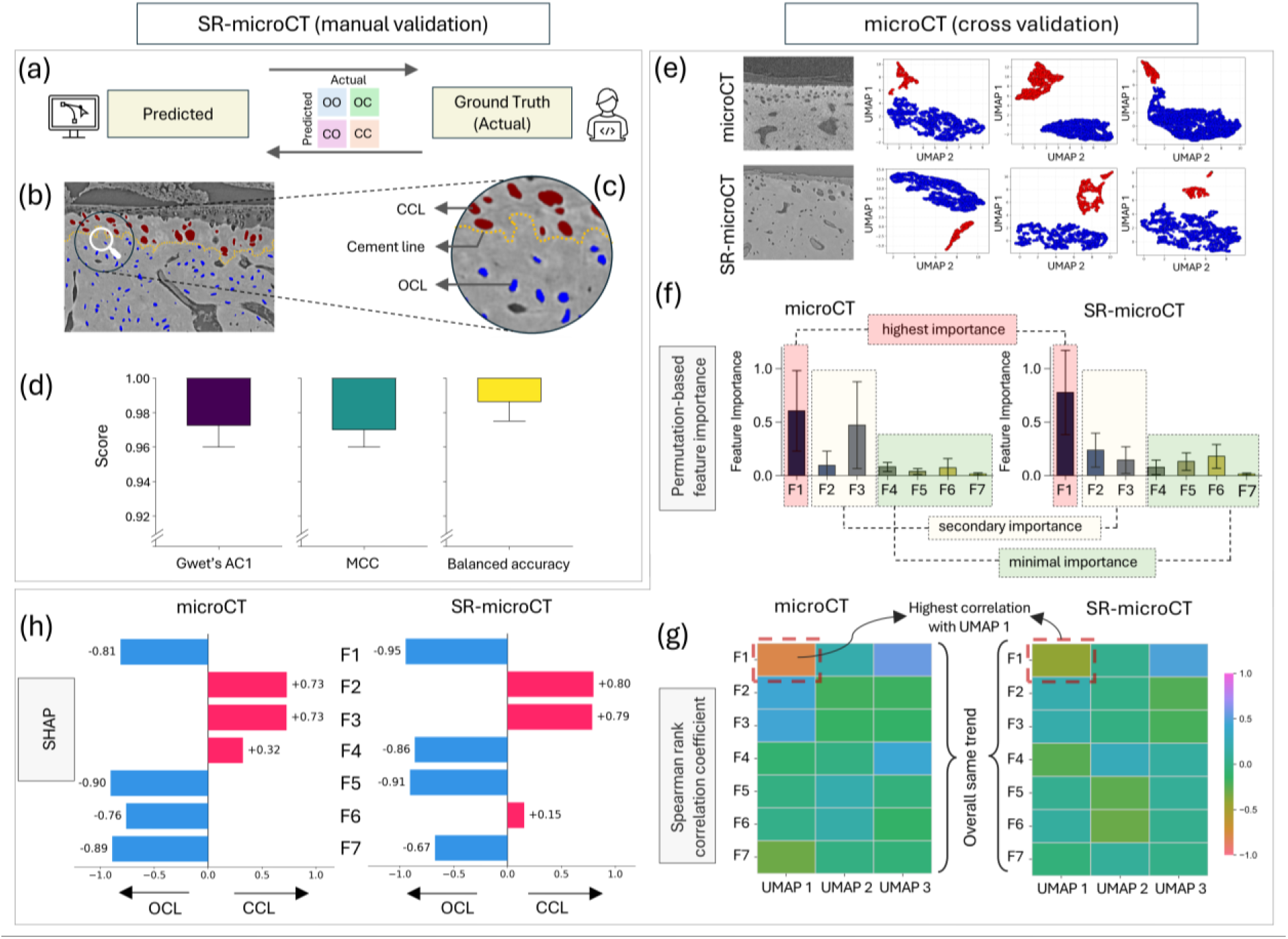
Classification validation in microCT and SR-microCT pipelines which produce equivalent classification structure. (a) Overview of the pipeline workflow, (b) starts with validation of classification in SR-microCT and validated with a trained observer, (c) back-tracked lacunae after classification with the pipeline (red= chondrocyte lacunae, blue=osteocyte lacunae, and yellow dashed line: cement line drawn manually for validation) and (d) Performance of the pipeline in SR-microCT data: boxplots of Gwet’s AC1, the Matthews correlation coefficient (MCC), and balanced accuracy (e) Cross-modality UMAP embeddings for microCT (top row) and SR-microCT (bottom row), three paired specimens each, at the optimal hyperparameters (n_neighbors = 5, min_dist = 0.01) (f) permutation-based feature importance for both modalities, with parameters labelled F1–F7, (g) Spearman rank correlation heatmaps (rows F1–F7; columns UMAP 1, UMAP 2, UMAP 3), and (h) SHAP diverging bar chart showing the Spearman rank correlation (ρ) between each parameters’ raw value and its SHAP contribution; bars extending right (positive ρ) indicate CCL tendency and bars extending left (negative ρ) indicate OCL tendency. Lc.D (F1) = lacunar distance, Lc.V (F2) = lacunar volume, Lc.SA (F3) = lacunar surface area, Lc.Φ (F4) = lacunar sphericity, Lc.E (F5) = lacunar elongation, Lc.F (F6) = lacunar flatness, Lc.Eq (F7) = lacunar equancy.

##### 2.2. Spearman Rank Correlation

Lc.D showed the strongest correlation with UMAP 1 in both modalities (SR-microCT: |r| = 0.41; microCT: |r| = 0.79) and a moderate correlation with UMAP 3 (SR-microCT: r = 0.48; microCT: r = 0.56). Lc.V and Lc.SA were the second and third strongest correlates of UMAP 1 in most datasets. In microCT, both reached moderate levels (r = 0.44 and 0.45), notably higher than in SR-microCT (r ≤ 0.34). Lc.E, Lc.F, Lc.Φ and Lc.Eq contributed selectively and at lower magnitudes. In both modalities, Lc. Eq showed the weakest and least consistent correlations (*Fig. 4g*).

##### 2.3. SHAP value analysis

Lc.D was the primary classification driver in both SR-microCT and microCT: higher Lc.D values consistently pushed lacunae toward OCL (SR-microCT: ρ = −0.81; microCT: ρ = −0.95). Lc.V and Lc.SA showed consistent CCL-associated tendencies in both modalities (SR-microCT: ρ = +0.73 for both; microCT: ρ = +0.80 and +0.79, respectively); lacunae with greater volume and surface area were pushed toward CCL. Lc.E and Lc.Eq were OCL-associated in both modalities, with higher values pushing toward OCL classification. Lc.Φ and Lc.F reversed direction between the two modalities. Altogether, five out of seven features showed consistent directional tendencies across modalities (*Fig. 4h*).

#### 3. Performance metrics (SR-microCT)

The 3D pipeline achieved near-perfect agreement with manual classification in SR-microCT. The mean Gwet’s AC1, MCC, and balanced accuracy were 0.99, 0.99, and 0.99, respectively, with median values of 1.00 for all three metrics. By the Landis and Koch thresholds, all mean AC1 values fall in the almost perfect agreement band (> 0.81)[29]. This indicates almost perfect classification of OCL and CCL [30], [31] (*Fig. 4d*).

### B. Validating the pipeline on microCT

#### 1. Cross-modality validation

Cross-modality validation was performed on the three DMM specimens scanned in both modalities, covering the same anatomical region. The remaining samples in each modality were used for single-modality analysis only. UMAP embeddings from microCT and SR-microCT data on the same three specimens produced qualitatively equivalent bimodal structure. HDBSCAN identified two clusters in both modalities at the same optimal hyperparameter configuration (*n_neighbors* = 5, *min_dist* = 0.01), which also yielded the highest silhouette scores in both modalities (microCT: 0.49; SR-microCT: 0.80). Silhouette scores were lower in microCT than in SR-microCT, and cluster boundaries were less compact.

Feature importance rankings were broadly consistent across modalities, with two notable differences. First, Lc.D remained the dominant parameter in both modalities but ranked first in only three of six microCT samples compared with four of six in SR-microCT. Second, Lc.V ranked second in SR-microCT whilst Lc.SA rose to second in microCT, a shift corroborated across all three importance analyses. Spearman correlations of Lc.V and Lc.SA with UMAP 1 were higher in microCT (r = 0.44–0.45) than in SR-microCT (r ≤ 0.34). Shape parameters remained secondary contributors in both modalities, and Lc.Eq was the lowest-ranked parameter in every sample. Five of seven parameters, Lc.D, Lc.V, Lc.SA, Lc.E, and Lc.Eq, maintained consistent SHAP directional tendencies across modalities.. Across all six comparison metrics, the microCT and SR-microCT pipelines produced equivalent classification structure on the same specimens, satisfying the pre-defined cross-modality validation criterion (*Fig. 4e-h*).

## DISCUSSION

This study presents an automated 3D pipeline to classify OCL and CCL in the osteochondral unit without separating the host tissues, subchondral bone and calcified articular cartilage, and without cement line tracing or labelled training data. Against SR-microCT, where manual classification is feasible, the pipeline achieved near-perfect agreement with a trained observer. Both in SR-microCT and microCT, the classification pipeline followed identical steps. The classification pipeline performed successfully in the microCT which is far more accessible than synchrotron infrastructure and does not need any complementary imaging modality, such as histology or backscattered electron imaging. This pipeline is therefore suitable for large-scale lacunar analysis in datasets where manual classification is impractical. Consistent performance across both healthy and post-traumatic osteoarthritis specimens demonstrates the pipeline’s applicability across biological conditions relevant to disease-focused research. This study also addressed a longstanding practical barrier to large-scale 3D characterisation of OCL and CCL together within osteochondral unit. Furthermore, the annotation-free design yields reproducible classifications independent of operator experience.

Compared with recent deep-learning approaches [13], [16] the present pipeline requires no labelled training data and no prior tissue separation, offering reproducibility, portability across datasets, and independence from annotation labour. On the specific task of OCL/CCL separation, near-perfect agreement with the manual gold standard was nonetheless achieved, demonstrating that supervised learning is not a prerequisite for accurate discrimination when the underlying feature structure is well-posed.

No single morphometric or positional parameter separated OCL from CCL reliably. Pairwise scatterplots confirm extensive overlap between the two populations across all individual measurements (*Fig. S4a*), consistent with prior reports that no single criterion, whether volume, shape, or orientation, draws a reliable boundary[13], [15]. This overlap is biologically expected at the osteochondral junction. Due to the complex nature of the cement line at the interface between calcified articular cartilage and subchondral bone, intermediate morphologies become more difficult to separate (*Fig. S4b; Fig. S11*). Parameters were selected to capture three complementary axes of biological variation: lacuna size descriptors (Lc.V and Lc.SA), three-dimensional shape descriptors (Lc.Φ, Lc.E, Lc.F, and Lc.Eq), and location within the mineralised compartment. Size descriptors quantify lacunar cavity dimensions and shape descriptors are derived from eigenvalue decomposition of the inertia tensor, capturing 3D anisotropy and geometric deviation from a spherical form. Lc.E, Lc.F, and Lc.Eq are three pairwise ratios of the squared semi-axis lengths from the same ellipsoidal fit and together they characterise the three-dimensional shape of the lacunar ellipsoid. The positional descriptor (Lc.D) encodes the depth of each lacunar centroid from the outer mineralised surface (*Fig. S1*). At the population level, OCL are smaller and more elongated, while CCL are larger, more spherical, and positioned closer to the joint surface [9], [18], [19].

Both the ground truth and the dominant feature, Lc.D, are based on positional reference. The observer classifies each lacuna based on its position relative to the cement line, whereas Lc.D is measured from the tidemark. The strong agreement between Lc.D-driven clustering and manual cement-line labelling reflects the anatomical continuity of these two positional references, not a circular definition of the labelling criterion. Lc.V and Lc.SA contributed consistently across both modalities, showing that morphometry adds classification signal beyond position alone. This multivariate basis is further supported by the cross-modality comparison. Consistent cluster structures, SHAP direction, and feature importance rankings were observed across majority of the features. These support the use of the 3D pipeline in microCT images, where the cement line is not visible. The shift from Lc.V to Lc.SA as the second-ranked feature in microCT does not alter the net classification outcome, as these two features are strongly correlated (r = 0.98) (*Fig. S3*).

The core classification framework developed for 3D images was extended to 2D images as an exploratory analysis to assess transferability beyond volumetric CT modalities. Unlike 3D morphometric descriptors, 2D descriptors reflect only the geometrical information available in a planar cross-section. The same lacuna can present different apparent morphology depending on the angle of sectioning. This is particularly significant for chondrocyte lacunae: a large chondron sectioned obliquely may appear as a small circular profile, which the algorithm may misread as an osteocyte lacuna. This geometric ambiguity inflates inter-class parameter overlap and occasionally produces multiple sub-clusters for the same class in 2D (*Fig. S12–S15*), whereas the 3D pipeline consistently yields one compact cluster per class. The lower performance is therefore a geometric consequence of dimensionality, not a limitation of the framework.

The present pipeline was developed and validated within a specific experimental context, and several directions remain open. Manual classification was performed by a single trained observer, ensuring a consistent segmentation style across all validation slices. The pipeline was validated on a single species and joint, and only for healthy and early-stage PTOA models. Future work should evaluate its performance across more diverse biological contexts where lacunar morphology may differ substantially from the present cohort. Silhouette scores were consistently lower in microCT than in SR-microCT, reflecting less compact cluster boundaries, and whether this difference affects classification robustness in specimens with more advanced morphological heterogeneity warrants further investigation. In this study, UMAP hyperparameters were optimised per sample rather than using a fixed configuration. Silhouette score was used to select the optimal hyperparameter combination, and this quantitative analysis provided the basis of the selection. Across all specimens, *n_neighbors* values of 5 and 10 combined with *min_dist* values of 0.01 and 0.05 produced well-separated clusters consistently. A supplementary analysis examining cluster separation under a fixed hyperparameter configuration across all specimens is provided in supplementary results (*Fig. S18*). However, for complete automation of the framework further investigation is required. When larger scan volumes are considered, lacuna morphology within each population may also vary, and characterising this intra-cluster heterogeneity remains an open question. Lastly, the pipeline performed consistently across healthy and diseased specimens, but how cluster structure and classification behaviour change with disease progression has not yet been examined.

## CONCLUSION

This study developed and validated a fully automated, annotation-free pipeline for classifying osteocyte and chondrocyte lacunae in the osteochondral unit without cement line tracing, tissue separation, or labelled training data. The pipeline achieved near-perfect agreement with manual classification in both healthy and post-traumatic osteoarthritis specimens. Classification was based on six morphometric parameters and a positional descriptor. Cross-modality equivalence between SR-microCT and microCT was established on paired specimens, extending the pipeline to a widely accessible modality. The framework transferred to two-dimensional imaging, with lower performance reflecting geometric ambiguity in planar sectioning rather than a failure of the classification logic. By removing the requirement for manual tissue delineation, this pipeline enables reproducible, large-scale characterisation of lacunar populations in the osteochondral unit and provides a foundation for future studies extending the pipeline beyond the present experimental model to diverse biological and clinical contexts.

## CODE AVAILABLE

The 3D and 2D classification pipeline used to perform all analyses in the paper, and the code and GUI are available at https://gitlab.kuleuven.be/u0161341/uloc. Raw SR-microCT and microCT reconstructed volumes and manually classified ground-truth labels are available from the corresponding author upon reasonable request.

## ACKNOWLEDGE

This work has received funding from the FWO and F.R.S.-FNRS under the Excellence of Science (EOS) programme (EOS No. 40007553), and KU Leuven Global PhD Partnerships with the University of Melbourne, Reference nr.: GPUM/22/019. The KU Leuven XCT Core Facility is acknowledged for the 3D image acquisition and processing tools (https://xct.kuleuven.be/). We would like to E. Pedrinazzi, L. Demeuldre and A. Malerba from University of Liège as well as J. Perrin and T. Weitkamp from Synchrotron SOLEIL for their help with SR-microCT. We would also like to thank Annunzia M Cagnoni, Han Liu and the animal facility staff of the University of Melbourne for their assistance with the animal study.

## Notes

### Competing Interest Statement

The authors have declared no competing interest.

