## Supplementary for "Unsupervised 3D classification and characterisation of osteocyte and chondrocyte lacunae in the murine osteochondral unit"

### Supplementary material

#### S1. Methods

##### Part A: 3D pipeline

##### A.1. Feature Normalisation

Prior to dimensionality reduction, each parameter was standardised to a z-score using the StandardScaler function from scikit-learn (v1.2.0). This normalisation ensured that each parameter contributed equally regardless of its original numerical range. Standardisation has been shown to improve both computational efficiency and classification accuracy (Pedregosa et al., 2011).

##### A.2. Feature extraction

To assess collinearity among input parameters prior to dimensionality reduction, Pearson correlation coefficients were computed between all pairwise parameter combinations for the 3D pipeline (Lc.V, Lc.SA, Lc.D, Lc.Φ, Lc.F, Lc.E, and Lc.Eq). Correlation values were visualised as symmetric heatmaps ($v_{min} = -1$, $v_{max} = +1$) to assess collinearity before dimensionality reduction. This preliminary analysis identified potential parameter redundancy and informed downstream analyses.

##### A.3. Positional parameter: lacunae distance (Lc.D)

In addition to the six morphological parameters, a positional parameter named lacuna distance (Lc.D), was calculated for each lacuna. From each CT slice (SR-microCT and microCT) mineralised tissue was segmented from the background. The outer surface curvature of the segmented region was identified as the joint outline using the "Outline" command in ImageJ. The topography of this outline approximately follows the tibial plateau as well as the cement line, providing a tissue-agnostic positional reference that does not require explicit cement line delineation. The Euclidean distance from each lacuna centroid to this outline was measured and registered as Lc.D (Fig. S1). Since microCT and SR-microCT resolve only mineralised tissue, all segmented lacunae (including OCL and CCL) reside on one side of the outline, ensuring that Lc.D captures the depth of each lacuna within the mineralised compartment. Lacunae with Lc.D values between 0 and 10 μm were excluded from all analyses to eliminate edge artefacts and partially segmented structures near the plateau surface.

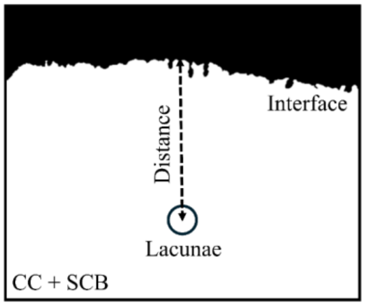


Fig. S1. *Schematic definition of the lacuna distance* parameter. *(lacunar distance = Lc.D ; CC = calcified cartilage ; SCB = subchondral bone; interface = tidemark)*

##### A.4. Dimensionality Reduction Techniques: PCA, t-SNE and UMAP

Three dimensionality reduction techniques, Principal Component Analysis (PCA), t-Distributed Stochastic Neighbour Embedding (t-SNE) and Uniform Manifold Approximation and Projection (UMAP), were evaluated to identify the approach best suited to separate OCL and CCL populations.

*Principal Component Analysis (PCA)* is a linear technique that transforms the standardised data into a set of orthogonal principal components capturing maximum variance. Three principal components (n_components = 3) were computed for each dataset. Pairwise 2D projections (PC1 vs PC2, PC1 vs PC3, and PC2 vs PC3) were visualised to assess the linear separability of lacunar populations.
t-Distributed Stochastic Neighbour Embedding (t-SNE) is a nonlinear technique that prioritises the similarities between neighbouring data points and preserves local structure in the low-dimensional embedding (van der Maaten and Hinton, 2008). Three-dimensional t-SNE embeddings (n_components = 3) were computed at three perplexity values to explore sensitivity across different structural perplexity scales: 10, 50, and 100 (Kobak and Berens, 2019). All t-SNE computations used random_state = 200 for reproducibility. The 2D projections of the three-dimensional embeddings were visualised for each perplexity setting to assess cluster formation qualitatively. Each technique was applied independently to the standardised parameter matrix of each sample, and the resulting low-dimensional embeddings, i.e. the reduced-coordinate space onto which every lacuna is projected, were qualitatively assessed for cluster separability. Cluster separability for each technique was optimised before the most suitable technique was selected for the classification pipeline. Selection criteria were: (i) clarity of cluster boundaries, (ii) distinctiveness of cluster separation, and (iii) consistency of clustering across different parameter subsets.

**Commented [pi1]:** Already providing findings here?

##### A.5. Scalability Analysis

To assess pipeline robustness across field-of-view (FOV), the 3D pipeline was independently applied to SR-microCT data at three scales: the original FOV (1×), twice the original FOV (2×), and three times the original FOV (3×). A constant depth of 195 µm was maintained across all three scales. The same three scales were subsequently applied to the 2D pipeline, where the FOV was extended by increasing the width of the imaged region (1×, 2×, and 3×) (Fig. S2).

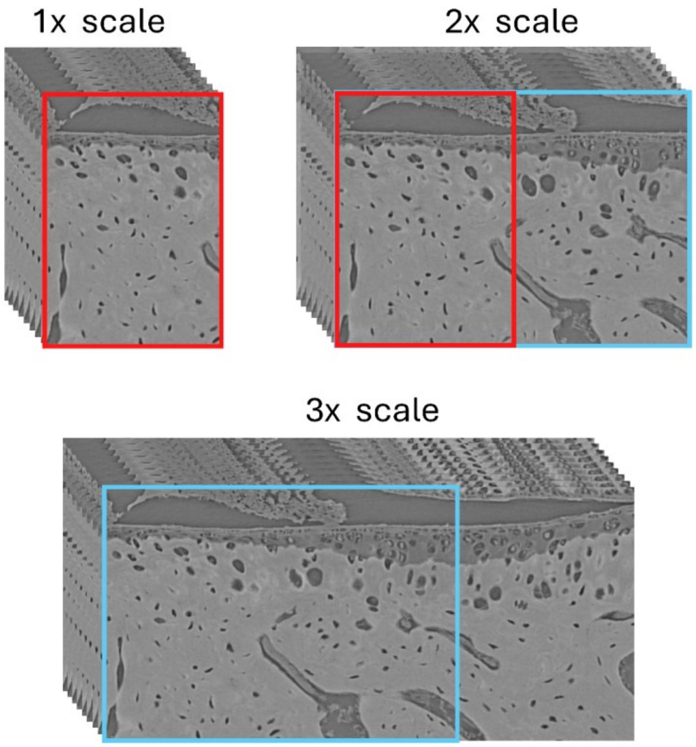


Fig. S2. *Field-of-view (FOV) scales used in the scalability analysis. Three stacked SR-microCT image blocks correspond to the 1×, 2×, and 3× scales. (3x is three times the 1x field of view; and 2x is twice the 1x field of view)*

Performance metrics (balanced accuracy, Gwet's AC1, Matthews Correlation Coefficient) and cluster counts were recorded at each scale and compared against manual counts to assess whether pipeline performance remained consistent as the volume or area of analysis increased. This scalability analysis evaluates whether the embedding space and clustering decisions are robust to variations in sampling region size.

##### A.6. Performance Metrics

*Gwet's AC1*

Gwet's AC1 was used as the primary inter-rater agreement metric, computed as:

$$AC1 = (P_o - P_e) / (1 - P_e) \qquad \text{Equation (1)}$$

where $P_o$ is the observed proportion of agreement, and $P_e$ is the expected agreement by chance. Agreement strength was interpreted using Landis & Koch's rule of thumb (Landis & Koch, 1977) and therefore, Gwet's AC1 values of 0-0.2, 0.21-0.4, 0.41-0.6, 0.61-0.8, and 0.81-1, indicate slight, fair, moderate, substantial, and almost perfect agreement levels respectively .

*Matthews Correlation Coefficient (MCC)*

The Matthews Correlation Coefficient (MCC) was computed to provide a balanced measure of classification quality that accounts for all four elements of the confusion matrix. MCC penalises the pipeline for errors in both classes (false positives and false negatives) and is not inflated by class imbalance. MCC ranges from +1 (perfect classification, which means the pipeline is in complete agreement with the manual classification) to −1 (perfect misclassification), with 0 indicating performance no better than random guessing (Chicco & Jurman, 2020; Zhu, 2020).

$$MCC = \frac{(TP \times TN) - (FN \times FP)}{\sqrt{(TP+FN)(TP+FP)(TN+FN)(TN+FP)}} \qquad \text{Equation (2)}$$

*Balanced Accuracy*

Balanced accuracy was computed as the arithmetic mean of per-class sensitivity:

$$Balanced\ accuracy = ½ \times \left(\frac{TP}{TP+FN} + \frac{TN}{TN+FP}\right) \qquad \text{Equation (3)}$$

This metric ensures that neither class is systematically favoured, which is critical when evaluating a pipeline on an inherently imbalanced dataset.

MCC is used as a balanced measure that accounts for all four confusion-matrix elements and is not inflated by class imbalance. Finally, balanced accuracy computed as the arithmetic mean of per-class sensitivity.

##### A.7. Cluster Interpretation via Back-Projection

Following HDBSCAN clustering in the low-dimensional UMAP space, the cluster labels assigned to each lacuna were back-projected onto the original three-dimensional image stacks to visualise the spatial distribution of OCL and CCL within the tissue. This was achieved using a 3D region-growing algorithm implemented in Python.

The centroid of each segmented lacuna in the volumetric image stack was identified. Each centroid was then used as a seed point from which the algorithm expanded outward into adjacent voxels using 26-connectivity (all 26 neighbours of each voxel in 3D space). The algorithm expanded into adjacent voxels with intensity greater than 200 or within a defined intensity threshold of the seed voxel (|intensity difference| < 30). These default threshold values were selected based on the intensity distribution of segmented lacunae in the reconstructed CT images.

This process reconstructed the spatial extent of each lacuna in the original image space and marked it according to its assigned cluster label. For the 2D pipeline, cluster labels were assigned directly by projecting each lacunar centroid onto the planar image and marking it with its corresponding HDBSCAN cluster label.

#### Part B: 2D Pipeline

##### B.1. Histological Sample Preparation

20 Safranin O-stained histological sections from four C57BL/6 mice (retired breeders, aged 7 months) were used to evaluate the performance of the 2D pipeline (five sections per mouse). This dataset was obtained as secondary data from a previous study(reference: https://doi.org/10.1002/jemt.20713). Proximal tibial condyles were dissected free from surrounding tissue and decalcified overnight in Morse's Solution (10% sodium citrate, 20% formic acid). Specimens were washed in running water for two minutes, then embedded in Tissue-Tek® O.C.T.™ compound (Sakura Finetek Europe, NL) and snap frozen in liquid nitrogen.

**Commented [AB2]:** I will include this after confirming the DOI with Kathryn

**Commented [GU3]:** The text you have is a direct copy of the 2009 paper, so probably worth including that reference here.

Specimens were oriented to allow sectioning in the sagittal plane starting laterally. Sections of 7 µm thickness were cut using a cryostat microtome (HM500 OV, Microm, DE) at −22°C until approximately midway through the lateral plateau. The last two sections were retained for analysis. Slides were stained with Safranin O, which binds to the proteoglycan content of cartilage and clearly delineates the boundary between cartilage and bone.

##### B.2. 2D Image processing and feature extraction

In the SR-microCT , the image processing is exactly identical to Method, section 2. In histology, the image processing for 2D segmentation followed an analogous workflow to the 3D pipeline, adapted for planar images.

In the histology, following Gaussian filtering (σ = 0.8), lacunae edges were detected using the "Edge Detection" command in ImageJ. Each closed outline was then segmented from the image, after which only the potential osteocyte and chondrocyte canalicular lacunae were retained through manual selection to exclude non-biological structures (e.g., cracks, imaging artefacts). The "Fill Holes" command was subsequently applied to delineate lacunar boundaries.

Once, the lacunae are extracted and successfully segmented in both SR-microCT and histology, two-dimensional morphological parameters were extracted for each segmented lacuna using the "Analyze Particles" command in ImageJ.

##### B.3. 2D morphometric parameters

For the 2D pipeline applied to SR-microCT slices and histological sections, six morphometric parameters (Table S1) were extracted:

Table S1. *Morphometric and positional parameters extracted per lacuna in the 2D pipeline.*

| Symbol | Parameter | Description |
|---|---|---|
| Lc.A | Lacuna cross-section area | Area of selected lacuna ($\mu m^2$) |
| Lc.P | Lacuna perimeter | The length of the outside boundary of the selected lacunae (µm) |
| Lc.C | Lacuna circularity | $4\pi \times \frac{[Area]}{[Perimeter]2}$ With a value of 1.0 indicating a perfect circle |
| Lc.AR | Lacuna aspect ratio | $\frac{[Major\ Axis]}{[Minor\ Axis]}$ The aspect ratio of the particle's fitted ellipse |
| Lc.S | Lacuna solidity | $\frac{[Area]}{[Convex\ area]}$ Convex area (Jarvis, 1973) |
| Lc.D | Distance of the lacunae centroid from the surface outline | Euclidean distance is calculated similar to Lc.D calculation in the 3D pipeline (µm) |

##### B.4. Weighted scaling in 2D pipeline

In the 2D classification pipeline, a weighted scaling approach was applied to selected morphometric parameters before dimensionality reduction and clustering. Specifically, higher weights were assigned to Lc.D (distance from surface) and Lc.A (lacunar area) to ensure these parameters had greater influence on the final clustering outcome. This weighting scheme was motivated by the observation that these two parameters are particularly discriminatory between OCL and CCL in 2D cross-sections, as CCL are typically larger and closer to the articular surface than OCL. Lc.D was calculated with a similar approach mentioned earlier.

##### B.5. Dimensional reduction techniques

The same UMAP and HDBSCAN parameters were applied as for the SR-microCT 2D pipeline (n_neighbors = 5, min_dist = 0.01, n_components = 3, min_samples = 9, min_cluster_size = 9).

The identical UMAP–HDBSCAN configuration was used across all three SR-microCT 2D scales (1×, 2×, 3×) and for the histology sections. Performance was evaluated against manual classification at 1× field-of-view (FOV) only for the histology dataset; SR-microCT 2D was evaluated at all three scales.

##### B.6. Performance Metrics

Classification performance of the 2D pipeline was quantified using the same three metrics defined in Section A.6: Gwet's AC1 (Equation 1), Matthews Correlation Coefficient (Equation 2), and balanced accuracy (Equation 3). For each 2D slice or histological section, cluster labels produced by the pipeline were compared with the manually assigned OCL/CCL labels to construct a per-slice confusion matrix, from which the three metrics were computed. Reported values are per-slice medians and means across all evaluated slices, matched to the corresponding modality and FOV scale (SR-microCT 2D: 1×, 2×, 3×; histology: 1× only). Interpretation of AC1 magnitudes followed the Landis & Koch bands described in A.6.

#### S2. RESULTS

##### Part A: 3D pipeline; SR-microCT

##### A.1. Lacunar Separation

Strong collinearity was observed between Lc.V and Lc.SA (r=0.98), while the positional parameter Lc.D remained independent of all morphological parameters. Lc.F vs Lc.E (r = −0.96), and Lc.F vs Lc.Eq (r=-0.87) showed a strong negative correlation. Lc.D showed weak correlation with every morphological parameter (|r| ≤ 0.26) (Fig. S3, S4).

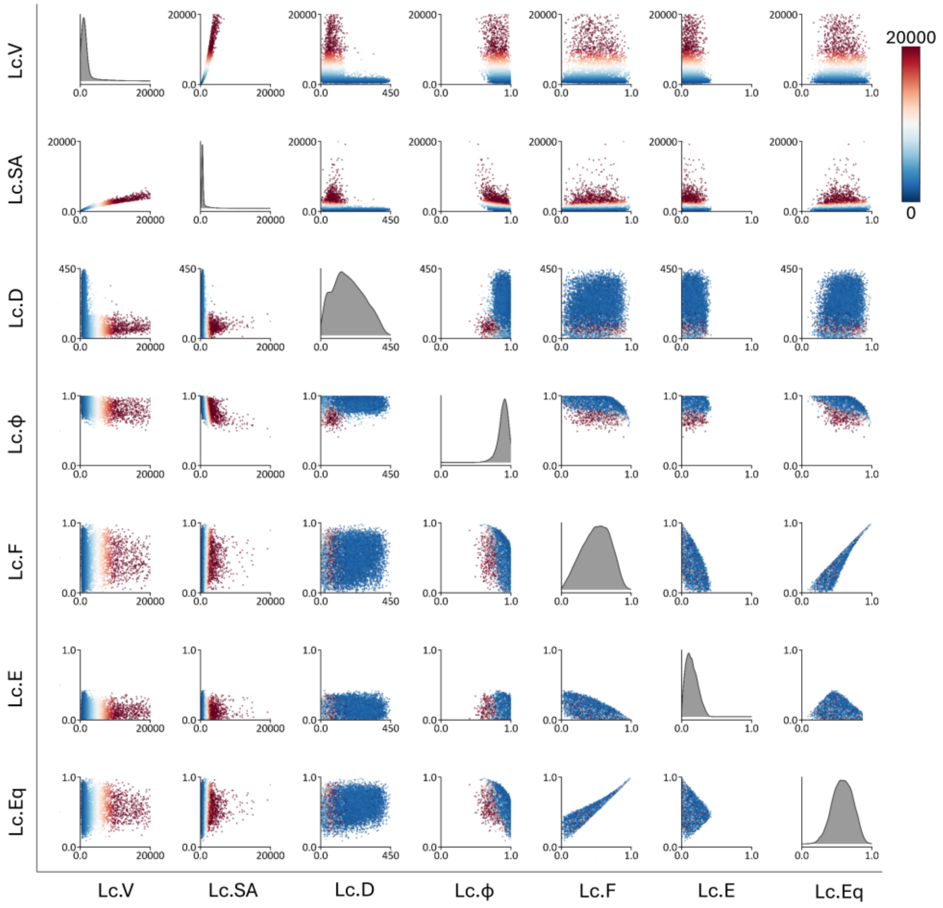


Fig. S3. *Feature correlation structure of the seven descriptors used in the 3D pipeline. (a) Pairwise scatterplot matrix of all* parameters*; off-diagonal entries show point density (blue, low; red, high)*

**Commented [Hv4]:** The panel is too small. Unreadable

**Commented [AB5R4]:** I would prefer to keep the scatter plot in the supplementary as it might take an entire page.

*and diagonal entries show the univariate distributions. (Lc.V = lacunar volume, Lc.SA = lacunar surface area, Lc.Φ = lacunar sphericity, Lc.E = lacunar elongation, Lc.F = lacunar flatness, Lc.Eq = lacunar equancy, Lc.D = lacunar distance)*

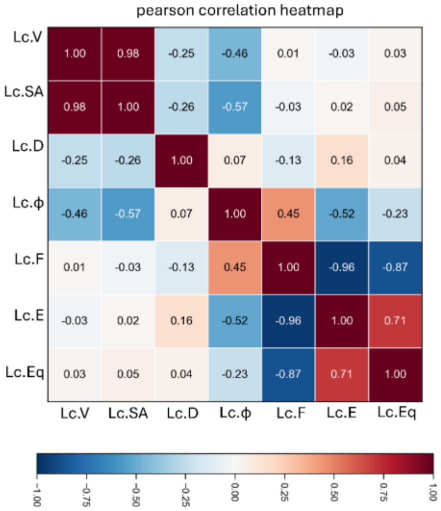


Fig. S4. *Pearson correlation heatmap of the same* parameters*, with the coefficient annotated in each cell on a scale from −1.00 (blue) to +1.00 (red). (Lc.V = lacunar volume, Lc.SA = lacunar surface area, Lc.Φ = lacunar sphericity, Lc.E = lacunar elongation, Lc.F = lacunar flatness, Lc.Eq = lacunar equancy, Lc.D = lacunar distance)*

##### A.2. Dimensionality Reduction Technique: PCA and t-SNE

Among the three techniques evaluated, UMAP produced the clearest bimodal separation between the two lacunae populations in both 3D and 2D classification. Performance was stable across scales, with a slight improvement at larger FOVs, attributable to the greater number of lacunae available for clustering.

- In PCA (Fig. S5), pairwise projections of the first three principal components showed substantial overlap between the two populations. No combination of components yielded a clear boundary.

- In t-SNE (Fig. S5), the embedding was fragmented into multiple clusters that were not distinctly separated at low perplexity (10). At intermediate perplexity (50), partial groupings were identified by HDBSCAN. At high perplexity (100), two broader regions emerged, but cluster edges remained diffuse. For perplexity values above 50, t-SNE showed reduced sensitivity and failed to capture the global structure (Kobak & Berens, 2019).

PCA yielded substantial overlap across all pairwise projections of the first three principal components, with no combination producing a clean boundary (Fig. S5). t-SNE produced fragmented and diffused cluster structure across all perplexity values tested (Fig. S5).

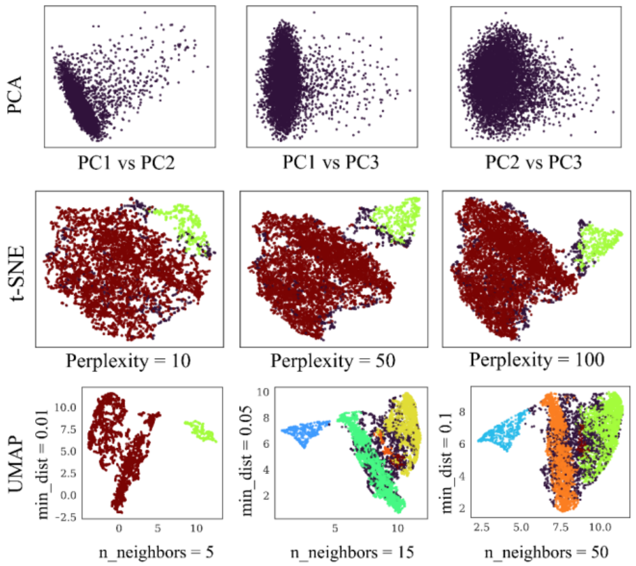


Fig. S5. *Comparison of dimensionality-reduction techniques and UMAP hyperparameter sweep for the 3D pipeline. (a) Embeddings from three techniques: principal component analysis (PCA; PC1 vs PC2, PC1 vs PC3, PC2 vs PC3), t-distributed stochastic neighbour embedding (t-SNE) at perplexities of 10, 50, and 100, and UMAP at n_neighbors of 5, 15, and 50 with min_dist of 0.01, 0.05, and 0.1,*

**Commented [Hv6]:** These are not shown, right?

**Commented [AB7R6]:** No, they are shown, and also mentioned in the image (Fig 3. (1))

##### A.3. UMAP Embedding Structure and Cluster Counts

At 1× and 2×, the highest mean silhouette scores were observed at n_neighbors = 5 and min_dist = 0.01 (1× = 0.45; 2× = 0.78), where HDBSCAN identified two clusters in each case (Fig. S6). The 3D pipeline consistently identified one OCL cluster and one CCL cluster at both scales (Fig. S6).

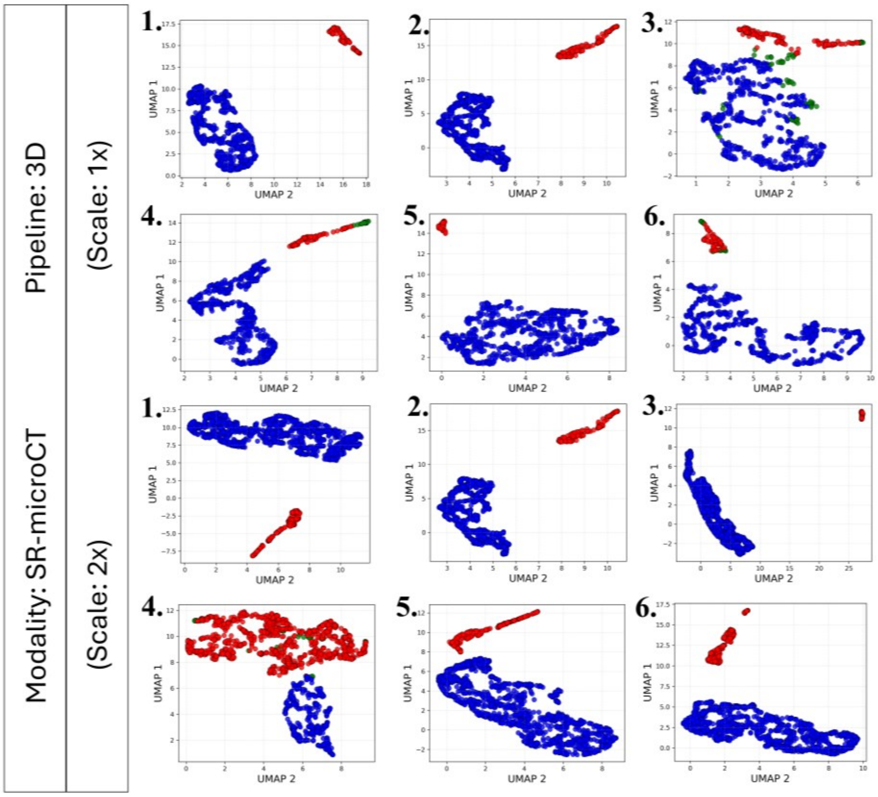


Fig. S6. *Per-specimen UMAP embeddings for the 3D SR-microCT pipeline at the 1× and 2× scales (UMAP 1 vs UMAP 2). (Blue = cluster 1; Red = cluster 2)*

#### A.4. Feature Importance

##### *Permutation-Based Feature Importance*

Lc.D ranked as the most important parameter in the majority of samples at both 1× and 2× scales, ranking first in four out of six samples at each scale (Fig. 5). In the remaining samples, Lc.V, Lc.E, or Lc.SA ranked first instead. The importance scores showed large variation across the six samples, as the standard deviation of Lc.D was approximately half of its mean at both scales. The mean importance of Lc.D was $0.83 \pm 0.45$ at 1× and $0.89 \pm 0.46$ at 2×. Lc.V had the second-highest mean at both scales ($0.26 \pm 0.17$ at 1× and $0.36 \pm 0.16$ at 2×). The shape parameters Lc.E, Lc.F, and Lc.φ scored below 0.18 at both scales. Lc.Eq was the lowest-ranked parameter at both scales (mean 0.01–0.02) (Fig. S7).

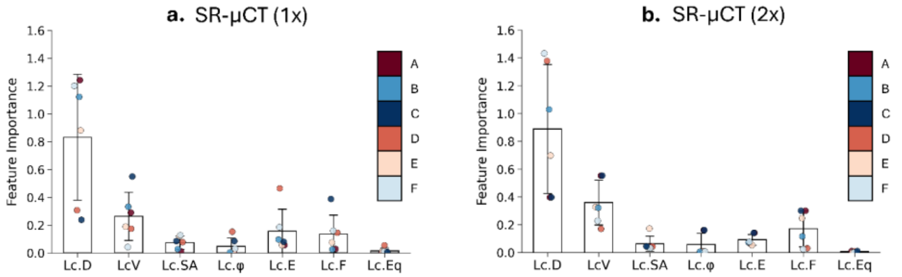


Fig. S7. *Permutation-based feature importance for the 3D SR-microCT pipeline at (a) 1× and (b) 2× scales. Lc.D = lacunar distance, Lc.V = lacunar volume, Lc.SA = lacunar surface area, Lc.Φ = lacunar sphericity, Lc.E = lacunar elongation, Lc.F = lacunar flatness, Lc.Eq = lacunar equancy.*

##### Spearman Rank Correlation

Lc.D showed the strongest correlation with UMAP 1 at both 1× and 2× scales (|r| = 0.72 and |r| = 0.40, respectively). At 1×, Lc.D also showed a moderate correlation with UMAP 3 (r = 0.47). Lc.V and Lc.SA were the second and third strongest correlates of UMAP 1 at both scales, remaining below r ≤ 0.34. Lc.F and Lc.Eq showed moderate correlations with UMAP 2 at 1× (r = 0.53 and 0.50), but this pattern did not persist at 2×. Lc.Eq showed the weakest and least consistent correlations at both scales (Fig. S8).

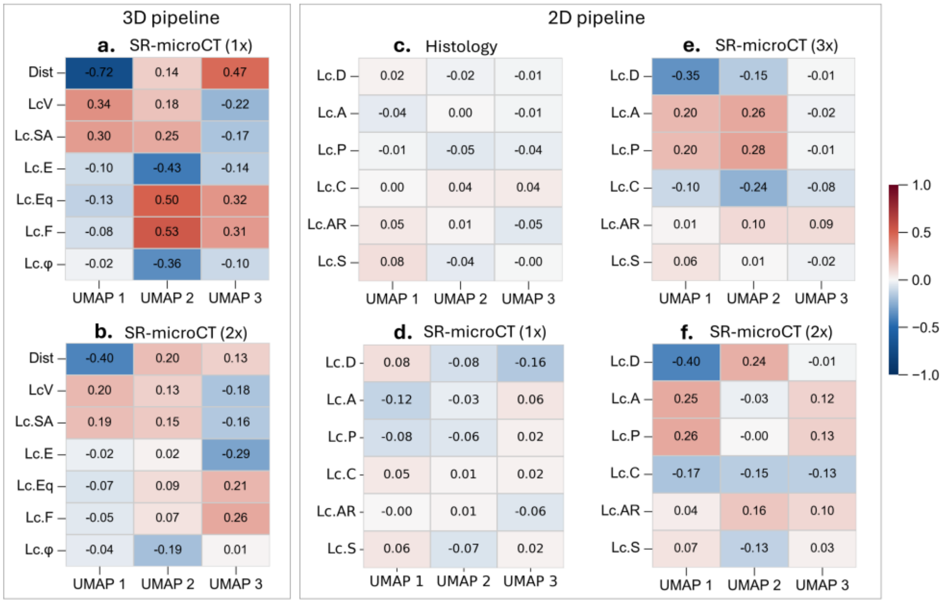

Fig. S8. *Spearman rank correlation heatmaps between morphological* parameters *and UMAP axes (UMAP 1, UMAP 2, UMAP 3), grouped by pipeline. For the 3D pipeline: (a) SR-microCT 1× and (b) SR-microCT 2×, with rows Lc.D, Lc.V, Lc.SA, Lc.E, Lc.Eq, Lc.F, and Lc.φ. For the 2D pipeline: (c) histology, (d) SR-microCT 1×, (e) SR-microCT 3×, and (f) SR-microCT 2×, with rows Lc.D, Lc.A, Lc.P, Lc.C, Lc.AR, and Lc.S. The coefficient is annotated in each cell on a scale from −1.0 (blue) to +1.0 (red). (Lc.D = lacunar distance, Lc.V = lacunar volume, Lc.SA = lacunar surface area, Lc.Φ = lacunar sphericity, Lc.E = lacunar elongation, Lc.F = lacunar flatness, Lc.Eq = lacunar equancy. (Lc.A = lacunar area; Lc.P = lacunar perimeter; Lc.D =lacunar distance; Lc.C =lacunar circularity; Lc.AR = lacunar aspect ratio; Lc.S = lacunar solidity).*

##### *Shapley Additive explanations (SHAP)*

Lc.D was the primary driver of classification decisions across all 3D SR-microCT datasets (1x and 2x), consistent with the permutation importance and Spearman correlation results. Lc.V and Lc.SA showed consistent secondary contributions across all datasets. Lacunae with low feature values (blue) were predominantly associated with negative SHAP, while higher values (red) tended toward positive SHAP (Fig. S9).

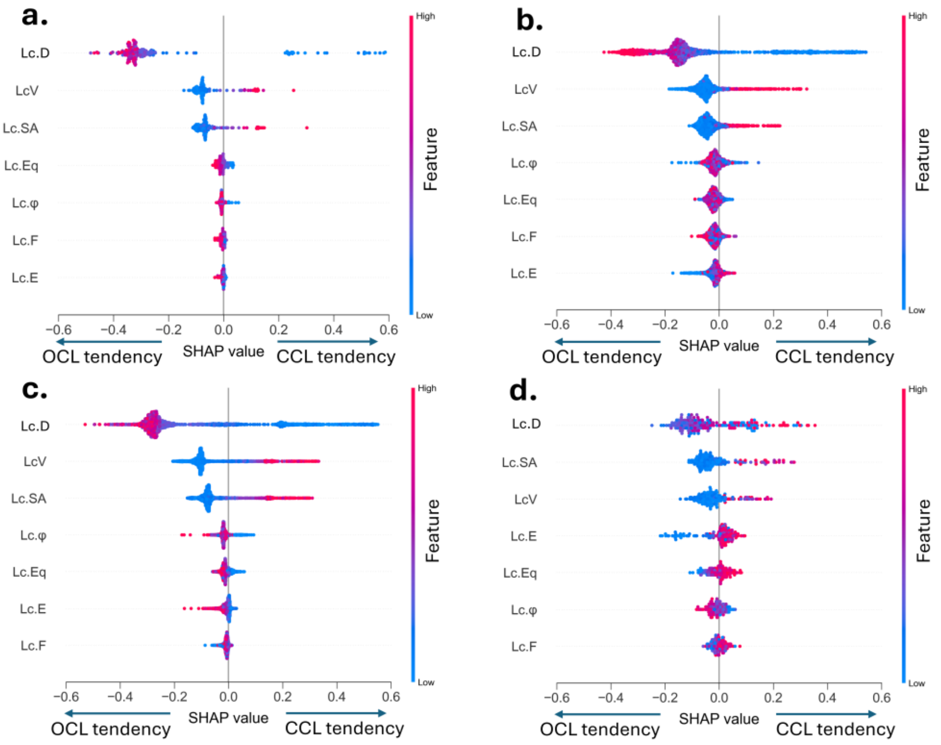

Fig. S9. *SHAP beeswarm plots for the 3D SR-microCT pipeline at (a) 1× (b) 2× (c) 3× and (d) microCT. Each point represents a single lacuna, the colour stands for a normalised parameter magnitude (red= high; blue= low); in the SHAP value axis, negative SHAP values indicating OCL tendency and positive values CCL tendency; (Lc.D = lacunar distance, Lc.V = lacunar volume, Lc.SA = lacunar surface area, Lc.Φ = lacunar sphericity, Lc.E = lacunar elongation, Lc.F = lacunar flatness, Lc.Eq = lacunar equancy).*

#### A.5. Number of Lacunae Analysed

Across 12 validation slices (6 specimens × 2 slices each), 563 lacunae were manually classified at 1× FOV (412 OCL, 151 CCL; total 46.9 ± 9.6, OCL 34.3 ± 9.4, CCL 12.6 ± 3.7 per slice) and 1,017 at 2× (728 OCL, 289 CCL; total 84.8 ± 18.7, OCL 60.7 ± 16.6, CCL 24.1 ± 9.1 per slice).

#### A.6. Performance

The 3D pipeline achieved near-perfect agreement with manual classification at both 1× and 2× FOV scales. The median values of Gwet's AC1, MCC, and balanced accuracy were 1.00 at both scales. The mean Gwet's AC1 was 0.95 at 1× and 0.98 at 2×. The mean MCC was 0.95 at 1× and 0.97 at 2×. The mean balanced accuracy was 0.98 at 1× and 0.99 at 2×. Performance improved slightly from 1× to 2×, attributable to the greater number of lacunae available for clustering at the larger scale (Fig. S10).

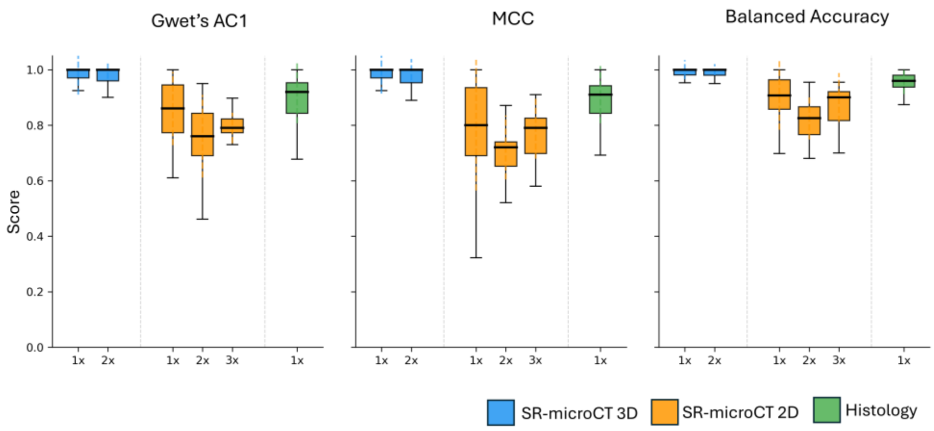


Fig. S10. *Classification performance across all pipelines and scales for (a) Gwet's AC1, (b) Matthews correlation coefficient (MCC), and (c) balanced accuracy, with scores (0–1). Boxplots are coloured as per imaging modality and pipeline: SR-microCT 3D (blue; 1×, 2×), SR-microCT 2D (orange; 1×, 2×, 3×), and histology (green; 1×).*

**Part B: 2D pipeline**

##### B.1. Lacunar Separation

In the 2D pipeline, a similar pattern to the 3D pipeline was observed. Lc.A and Lc.P were strongly correlated (r = 0.97). Lc.C and Lc.AR showed a negative correlation (r = −0.72). Lc.D remained weakly correlated with all shape parameters ($|r| \leq 0.40$). Pairwise scatterplots confirmed that no single parameter pair was able to separate the OCL and CCL populations cleanly, due to the extensive overlap of parameter distributions (Fig. S11).

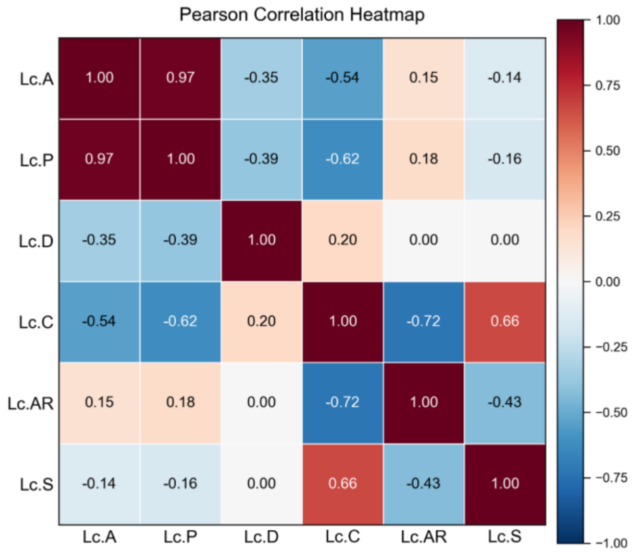


Fig. S11. *Pearson correlation heatmap of the six 2D morphological* parameter*s. The coefficient is annotated in each cell on a diverging scale from −1.00 (blue) to +1.00 (red). (Lc.A = lacunar area; Lc.P = lacunar perimeter; Lc.D =lacunar distance; Lc.C =lacunar circularity; Lc.AR = lacunar aspect ratio; Lc.S = lacunar solidity).*

##### B.2. Dimensionality Reduction Technique: PCA and t-SNE

Based on the qualitative comparison across dimensionality-reduction techniques described for the 3D pipeline (Result A.2), which showed identical trends when replicated on 2D descriptors, UMAP at the configuration n_neighbors = 5 and min_dist = 0.01 was selected for the 2D pipeline, with adjusted HDBSCAN hyperparameters (min_samples = 9, min_cluster_size = 9).

##### B.3. UMAP Embedding Structure and Cluster Counts

In the SR-microCT 2D pipeline, cluster counts were less stable across scales. Two OCL clusters appeared in one, three, and six slices at 1×, 2×, and 3×, respectively. For CCL, two clusters were identified in one, two, and two slices at 1×, 2×, and 3×, respectively, and three clusters appeared in one slice each at 2× and 3×. In the histology pipeline, one OCL cluster was identified in all 20 sections. Five sections produced two CCL sub-clusters rather than one. (Fig. S12-S14).

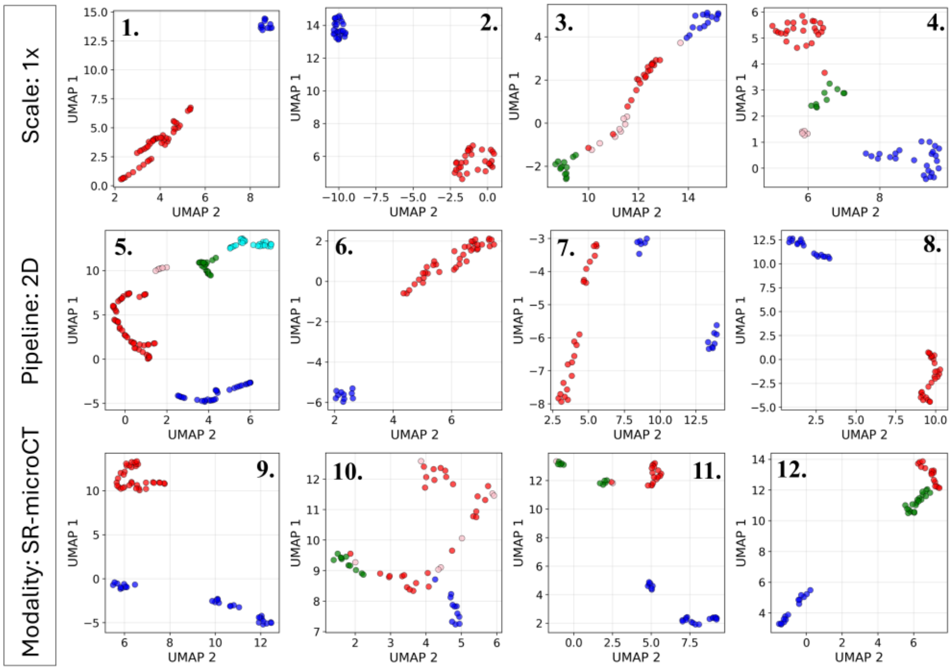


Fig. S12. *Per-specimen UMAP embeddings for the 2D SR-microCT pipeline at the 1× scales (UMAP 1 vs UMAP 2). (red = cluster 1; pink = cluster 2; blue = cluster 3; cyan = cluster 4; green = cluster 5)*

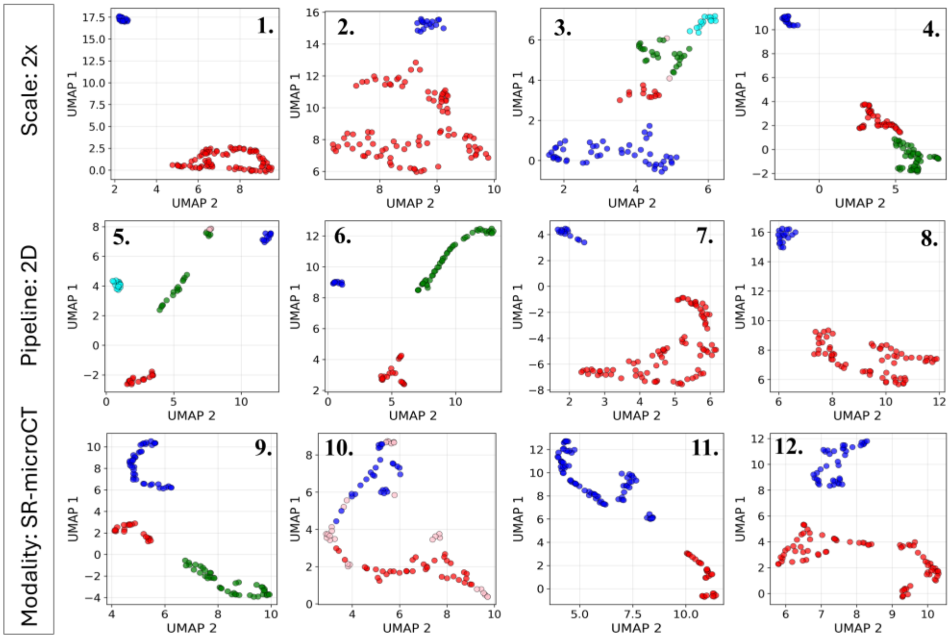


Fig. S13. *Per-specimen UMAP embeddings for the 2D SR-microCT pipeline at the 2× scales (UMAP 1 vs UMAP 2). (red = cluster 1; pink = cluster 2; blue = cluster 3; cyan = cluster 4; green = cluster 5)*

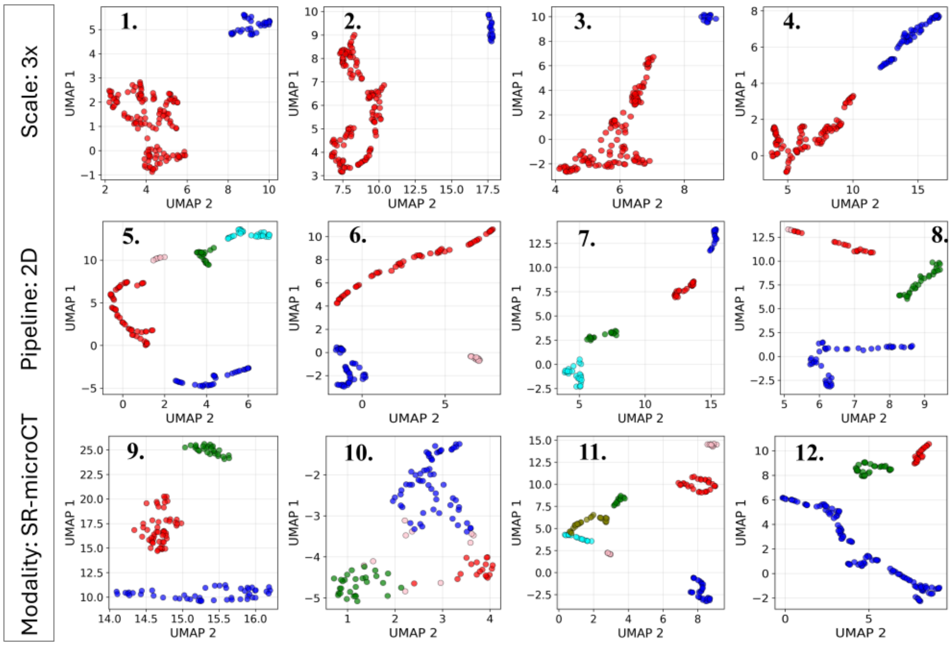


Fig. S14. *Per-specimen UMAP embeddings for the 2D SR-microCT pipeline at the 3× scales (UMAP 1 vs UMAP 2). (red = cluster 1; pink = cluster 2; blue = cluster 3; cyan = cluster 4; green = cluster 5)*

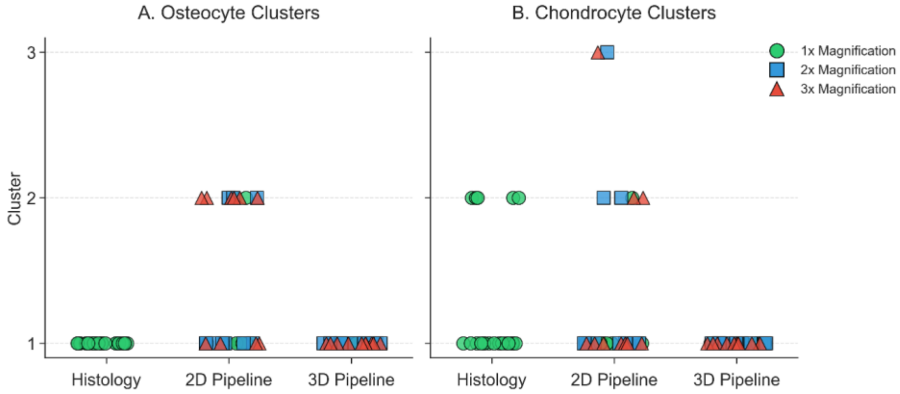


Fig. S15. *Summary of cluster counts across pipelines and modalities for (a) osteocyte clusters and (b) chondrocyte clusters, in each specimen; (x-axis = the cluster counts (1, 2, or 3) and y-axis = imaging modalities (histology, 2D pipeline, 3D pipeline)); the marker shape encodes*

*magnification (circle = 1×; square = 2×; triangle = 3×), and each marker represents one specimen (3D = VOIs and 2D = slices) or histology section (VOI = volume of interest).*

##### B.4. Feature Importance

*Permutation-Based Feature Importance*

In all three SR-microCT 2D scales, Lc.D had the highest mean importance (1.20 ± 0.63 at 1×, 1.00 ± 0.48 at 2×, and 1.07 ± 0.29 at 3×). The gap between Lc.D and all other parameters was wider in the 2D pipeline than in the 3D pipeline. All other parameters scored below 0.19 at every scale. Lc.A and Lc.P were the second and third-ranked parameters at 2× and 3×. Lc.S was the lowest-ranked parameter at all three scales.

In histology, no single parameter stood out. The mean importance of all six parameters fell between 0.15 and 0.22. Lc.S had the highest mean (0.22), followed by Lc.D, Lc.C, and Lc.AR at approximately 0.20 each (Fig. S16).

``

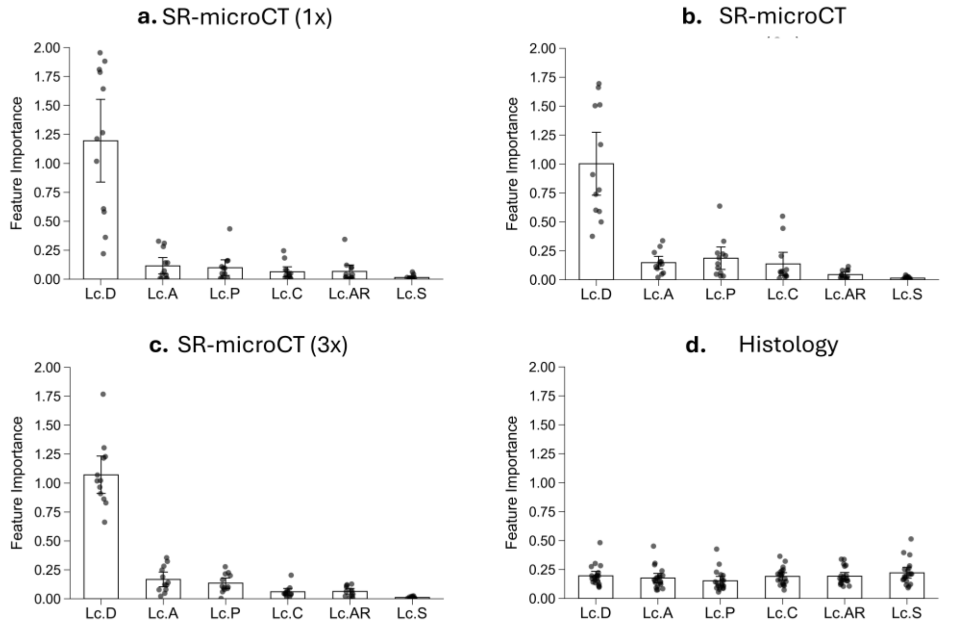


Fig. S16. *Permutation-based feature importance for the 2D pipeline across modalities: (a) SR-microCT 1×, (b) SR-microCT 2×, (c) SR-microCT 3×, and (d) histology. Each points represents each 2D slices of SR-microCT and histology. Lc.A = lacunar area, Lc.P = lacunar perimeter, Lc.D = lacunar distance, Lc.C = lacunar circularity, Lc.AR = lacunar aspect ratio, Lc.S = lacunar solidity*

##### *Spearman Rank Correlation*

Overall, correlations in the SR-microCT 2D pipeline were weaker than in the 3D pipeline across all scales. No parameter reached $|r| > 0.40$. Lc.D remained the strongest correlate of UMAP 1 at 2× ($r = -0.40$) and 3× ($r = -0.35$). At 1×, all correlations including Lc.D were weak ($|r| \leq 0.16$), with Lc.D showing only a marginal association with UMAP 3 ($r = -0.16$). This scale-dependent pattern mirrors the permutation importance results, where Lc.D dominated at all scales but with greater expression at coarser resolutions (Fig. S8). In histology, all correlations were very weak ($|r| \leq 0.08$), with no parameter showing a meaningful association with any UMAP axis (Fig. S8).

#### B.5. Performance

##### *SR-microCT*

The 2D SR-microCT pipeline showed lower agreement with manual classification than the 3D pipeline across all scales. At 1× FOV, the median Gwet's AC1, MCC, and balanced accuracy were 0.86, 0.80, and 0.91, with corresponding mean values of 0.84, 0.76, and 0.88. At 2× FOV, the median Gwet's AC1, MCC, and balanced accuracy decreased to 0.76, 0.72, and 0.83, and the mean decreased to 0.74, 0.70, and 0.82. At 3× FOV, performance recovered slightly relative to 2×; the median values were 0.79, 0.79, and 0.90, and the mean values were 0.80, 0.76, and 0.86 for Gwet's AC1, MCC, and balanced accuracy, respectively. Variability between samples was greater than in the 3D pipeline at all three scales. The 3× FOV showed the narrowest interquartile range, indicating the greatest consistency across specimens, while 1× showed the highest mean accuracy across all three metrics (Fig. S10).

##### *Histology Pipeline*

The histology pipeline outperformed the 2D SR-microCT pipeline. Across 20 sections, the median Gwet's AC1, MCC, and balanced accuracy were 0.92, 0.91, and 0.96, and the mean values were 0.88, 0.88, and 0.95. The interquartile range was 0.84–0.95 for AC1, 0.84–0.94 for MCC, and 0.94–0.98 for balanced accuracy, indicating a narrow spread and consistent performance across most sections. The histology pipeline was evaluated at 1× FOV only (Fig. S10) (Fig. S17).

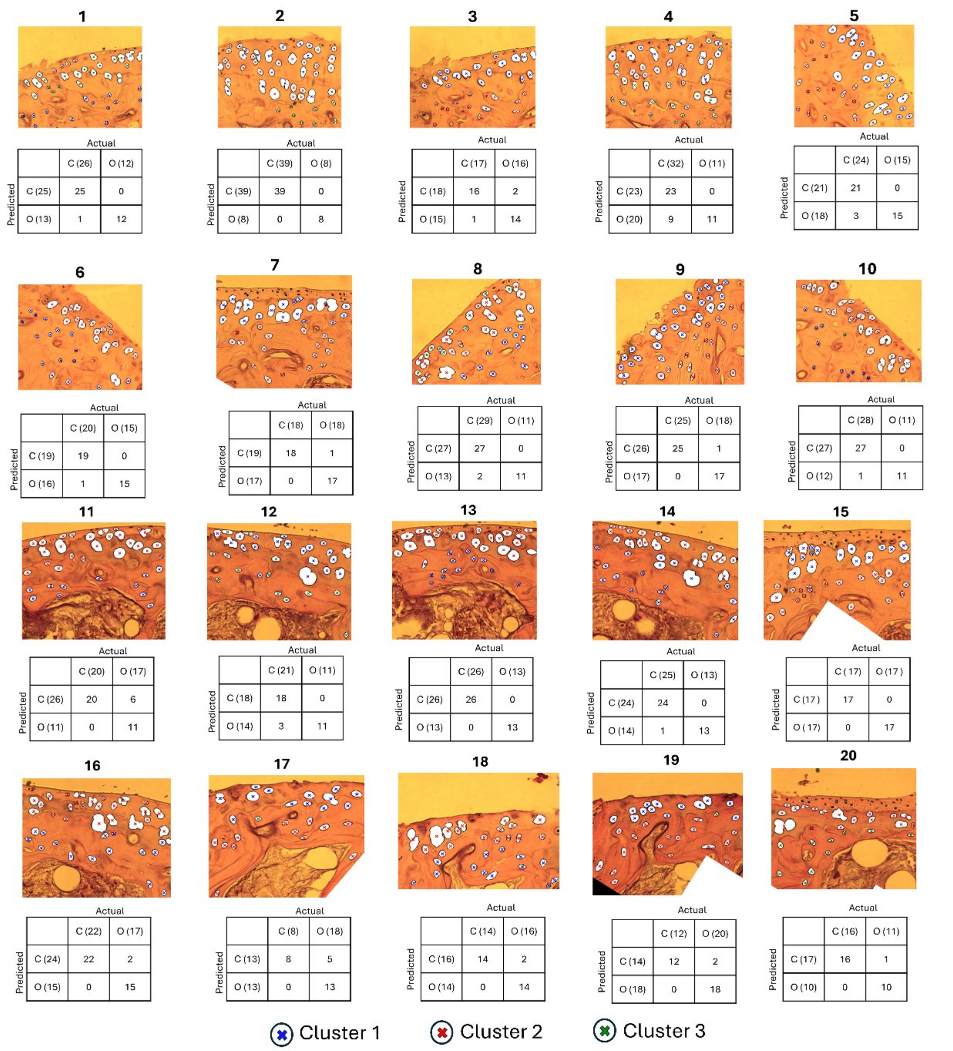


Fig. S17. *Per-section histology classification results for all 20 sections. Each table represents the confusion matrix of the particular specimen (rows: actual number of chondrocyte and osteocyte lacuna as per human observer; columns: predicted number of chondrocyte and osteocyte lacuna as per classification pipeline; (C = chondrocyte lacunae ; O: osteocyte lacunae). The marker style represents a cross and colours represent a single cluster with a single colour.*

#### B.6. Number of Lacunae Analysed

For histology, 727 lacunae were manually classified across 20 sections (288 OCL, 439 CCL; total 36.4 ± 5.3, OCL 14.4 ± 3.3, CCL 21.9 ± 7.3 per section).

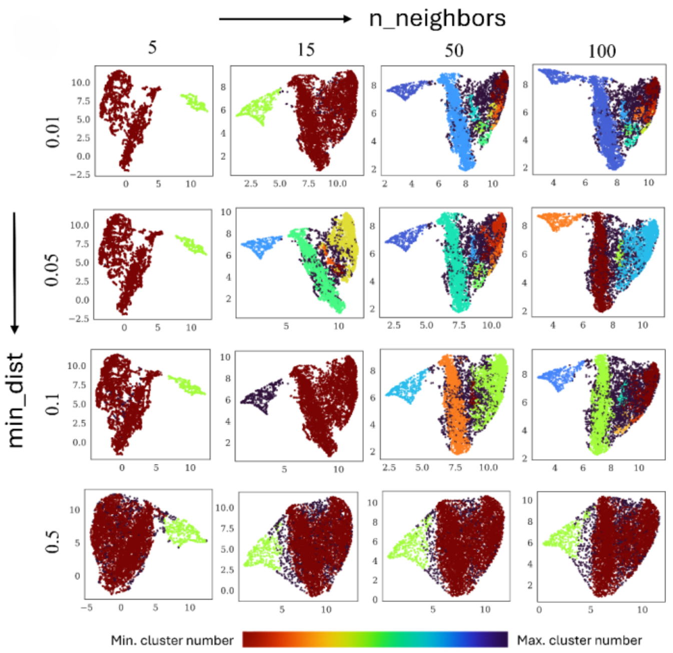


*Fig. S18. A projected UMAP sweep illustrating a representative subset of 16 configurations (columns: n_neighbors = 5, 15, 50, 100; rows: min_dist = 0.01, 0.05, 0.1, 0.5); the full analysis used 55 configurations as described in Methods A.3.Cluster number are based on HDBSCAN.*
